# The robustness of phylogenetic diversity indices to extinctions

**DOI:** 10.1101/2022.06.28.498028

**Authors:** Kerry Manson

## Abstract

Phylogenetic diversity indices provide a formal way to apportion evolutionary history amongst living species. Understanding the properties of these measures is key to determining their applicability in conservation biology settings. In this work, we investigate some questions posed in a recent paper by Fischer, Francis & Wicke appearing in *Systematic Biology* (Vol. 72(3), 2023). In that paper, it is shown that under certain extinction scenarios, the ranking of the surviving species by their Fair Proportion index scores may be the complete reverse of their ranking beforehand. Our main results here show that this behaviour extends to a large class of phylogenetic diversity indices, including the Equal-Splits index. We also provide a necessary condition for reversals of Fair Proportion rankings to occur on phylogenetic trees whose edge lengths obey the ultrametric constraint. Specific examples of rooted phylogenetic trees displaying these behaviours are given and the impact of our results on the use of phylogenetic diversity indices more generally is discussed.

## 1 Introduction

Each species on earth is the product of some evolutionary history, both unique to itself and shared with other species. Phylogenetic diversity indices are a family of measures that quantify this history on a species-by-species basis. They do so by assigning to each species a numerical score that aims to indicate that species’ contribution to biodiversity. One characteristic of diversity indices is that they calculate this contribution based on species’ positions in a rooted phylogenetic tree. This is in contrast to other approaches such as measuring the mean ‘patristic distance’ to other species or the shortest such distance, the ‘pendant edge length’ approach [1]. Moreover, unlike distance-based methods which measure the difference between species and their close relatives, diversity indices use phylogenetic information going right back to the root of the phylogeny under investigation.

Diversity indices and the scores they provide have found practical application in conservation biology. In particular, they have been used to suggest prioritisation rankings for biodiversity conservation, often in conjunction with other measures. For example, see [2–5] and other work by the EDGE of Existence programme [6]. One aim of using diversity indices is to move beyond conservation that overwhelmingly protects the most charismatic species at the expense of others [7]. The use of these measures can inform the distribution of conservation resources, prioritising those programmes that spread benefits and protection more widely, and properly reflect the full breadth of biodiversity.

That said, conservation efforts take place at a time heavily impacted by extinctions of species [8] and these extinction events impact the very measures that conservation biologists use to try to prevent them. Figuratively speaking, extinctions remove branches from the ‘tree of life’, thereby altering the phylogenetic tree structure on which these measures are based. For a conservation programme based on diversity indices, each extinction event necessitates the recalculation of diversity index scores for surviving species. This may lead to changes in relative rankings among species. Since conservation programmes often employ large amounts of scarce resources [9] and require community buy-in [10], it is useful to know if and when extinction events could lead to recalculations that give markedly different results. Large changes in priorities, and any subsequent reallocation of resources that followed, would not be conducive to the long-term planning required of many conservation programmes. Ideally, phylogenetic diversity indices would not only give informative prioritisation rankings but also be ‘robust’, that is, not particularly sensitive to changes caused by extinction.

Two diversity index methods have proven to be most popular, essentially to the exclusion of other approaches in this area. These are the Fair Proportion (FP) [2, 11] and Equal-Splits (ES) [1, 11] indices. Nonetheless, it is possible to define many other diversity indices that follow our definition (see p. 5) and we give some examples of these in Section 2.1. But in current practice only FP and ES are diversity indices of any significance. A third diversity index based on the Shapley value from co-operative game theory [12] has also been considered. Yet it was subsequently shown that on rooted phylogenetic trees the Shapley value is identical to the Fair Proportion index [13]. Other evolutionary isolation measures exist, such as those evaluated in [1] and a measure based on another co-operative game approach, the Banzhaf index (discussed in Supplementary material). However, none of these measures are phylogenetic diversity indices by the definition used here, largely ignoring edge lengths altogether. For this reason they will be set aside in the present discussion.

A recent paper by Fischer, Francis and Wicke [14] assesses the robustness of the Fair Proportion index on rooted phylogenetic trees. Those authors showed that species rankings under the FP index can change markedly following certain extinction events. Furthermore, “for each phylogenetic tree, there are edge lengths such that the extinction of one leaf per cherry completely reverses the ranking” [14, p. 2]. We call such an outcome a ‘ranking reversal’ induced by a set of extinctions. Their results raise concerns about the appropriateness of the Fair Proportion index and thus there would seem to be merit in developing an alternative. Some recent theoretical work has begun to look at properties of diversity indices in an abstract sense, with a view to evaluating their effectiveness as measures of species-level diversity [15, 16]. One motivating factor is that by taking a properties-first approach it may be that we can discover useful diversity index measures beyond FP and ES that do not share their flaws. However, our main results here (Theorems 5 and 9) show that a lack of robustness is exhibited by a large class of diversity indices. Moreover, we argue that those diversity indices outside this class have an unrealistic basis and thus any biologically reasonable diversity index used as an alternative to FP will not avoid the robustness problem. For this reason we do not claim that any alternative diversity index is unambiguously better than FP or ES for diversity index applications.

The FP index was a natural choice for [14] to begin the study of robustness of diversity indices. Given its relative importance, we also investigate the ES index specifically. The ES index generally requires more species to go extinct to induce a ranking reversal than FP does. This number may still be small though, and we give an example of a ranking reversal under ES caused by the extinction of just two species. Following Fischer and colleagues, we initially place no constraint on the size of edge lengths required to obtain such reversals. The examples contained in the original paper, the constructive methods appearing in the proofs of their Theorem 2 [14, Supp. Mat. pp. 2-15] and Theorems 5 and 9 here all tend to place the leaves at quite varied distances from the root vertex. This leads to questions about whether similar results can be obtained when edge lengths obey an ultrametric constraint. In particular:

> “if we restrict the analysis to ultrametric trees where all leaves have the same distance to the root, what are the worst-case scenarios in this setting?” [14, p. 5]

By “worst-case” those authors refer to a combination of edge lengths and extinctions that re-orders the FP index score ranking as much as possible. In Section 5 we outline some necessary conditions for a reversal of FP index scores in the ultrametric context. We then present examples of ultrametric rooted phylogenetic trees for which FP index score rankings are completely reversible. This answers the question above by showing that the “worst-case scenario” on an ultrametric phylogenetic tree is as bad as possible.

The rest of this paper is organised as follows. We begin with a section of preliminary definitions and notation before describing a handful of diversity indices besides FP and ES. The following two sections contain our main results: that two large classes of diversity index, called ‘non-rigid interior’ and ‘rigid interior’ indices respectively, are not robust in the sense described above. Section 5 focusses on robustness under an ultrametric constraint on edge lengths and is followed by a short final section of concluding remarks.

## 2 Preliminaries

Let *X* be a non-empty set of taxa (e.g. species), with |*X*| = *n*. A *rooted phylogenetic X-tree* is a rooted tree *T* = (*V, E*), where *X* is the set of leaves, and all edges are directed away from a distinguished root vertex *ρ*. An *interior* vertex of *T* is any vertex that is not a leaf. A rooted phylogenetic tree is called *binary* if every interior vertex has out-degree 2. All edges drawn in this paper will be directed down the page.

Let *P* (*T* ; *ρ, υ*) be the unique path in *T* from the root *ρ* to *υ* ∈ *V* (*T*). For any edge *e* ∈ *E*(*T*), we write *x* ∈ *c*_*T*_ (*e*) if *x* ∈ *X* and *P* (*T* ; *ρ, x*) includes *e*. That is, *c*_*T*_ (*e*) is the set (*cluster*) of leaves descended from the terminal vertex of *e*. For *υ* ∈ *V* (*T*), we also write *x* ∈ *c*_*T*_ (*υ*) if *x* ∈ *X* and *P* (*T* ; *ρ, x*) includes *υ*. If the (directed) edge (*u, υ*) appears in a phylogenetic tree *T*, we say that *u* is the *parent* of *υ*. If two distinct leaves *x*_1_ and *x*_2_ in *X* have the same parent vertex *υ* and no other vertex in *V* (*T*) has *υ* as a parent then we call *{x*_1_, *x*_2_*}* a *cherry*. A rooted binary phylogenetic tree with exactly one cherry is called a *caterpillar* tree. If a set of *m* ≥ 2 leaves *Y* ⊆ *X* all have the common parent vertex *v* and no other vertex in *V* (*T*) has *v* as a parent then we call *Y* an *m-cherry*.

The term ‘pendant’ is used in two related senses in this paper. First, a *pendant edge* is an edge whose terminal vertex is a leaf. Second, a *pendant* subtree of *T* is any subtree that does not contain the root vertex and can be a connected component of a graph formed from *T* by the deletion of exactly one edge. We write *P*_*e*_ for the pendant subtree that would be formed from the deletion of edge *e*. A rooted phylogenetic tree has a number of *maximal* pendant subtrees equal to the out-degree of the root, each formed by the deletion of an edge incident with the root.

We denote the two maximal pendant subtrees of a rooted binary phylogenetic tree *T* by *T*_*a*_ and *T*_*b*_. Further subtrees, in the nonbinary case, shall be denoted similarly. The tree shape of *T* may be expressed in terms its maximal pendant subtrees by writing *T* = (*T*_*a*_, *T*_*b*_, …). We may extend this notation to non-root non-leaf vertices of *T*, writing *T*_*a*_(*υ*), *T*_*b*_(*υ*), and so on, for the maximal pendant subtrees contained within the pendant subtree rooted at vertex *υ*. The edge connecting the root vertex to *T*_*a*_ will be labelled *a*, and the set of leaves in *T*_*a*_ will be denoted *X*_*a*_, with |*X*_*a*_| = *n*_*a*_. Parallel notation applies to the other maximal pendant subtrees of *T*. Figure 1 illustrates this notation for the simplest case, where *ρ* has out-degree 2.

**Fig. 1:**
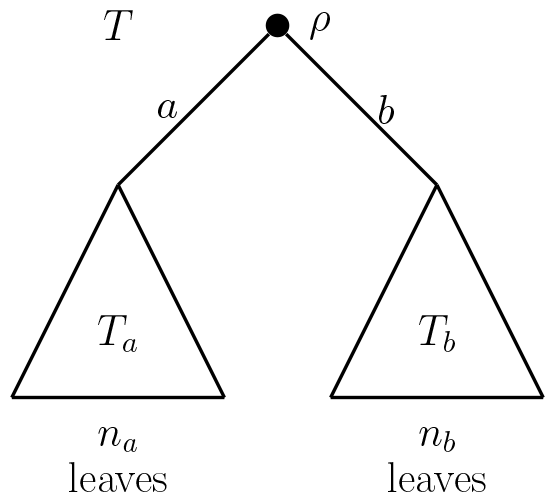
Schematic diagram showing the notation for a rooted phylogenetic tree *T* and its maximal pendant subtrees *T*_*a*_ and *T*_*b*_, which are descended from the root vertex *ρ* via edges *a* and *b*.

The edges of rooted phylogenetic trees in this paper are positively weighted. Let *T* be a rooted phylogenetic *X*-tree and let *𝓁* : *E*(*T*) → ℝ^*>*0^ be a function that assigns a positive real-valued length *𝓁* (*e*) to each edge *e* ∈ *E*(*T*). Suppose that *u, υ ∈ V* (*T*) are two vertices of *T* connected by a directed path from *u* to *υ*. Then the *distance from u to υ*, denoted *d*(*u, υ*), is the sum of the lengths of the edges in this path. If *𝓁* is such that for every two distinct leaves *x* and *y* we have *d*(*ρ, x*) = *d*(*ρ, y*), we say that *𝓁* satisfies the *ultrametric* condition.

A (phylogenetic) *diversity index* on a rooted phylogenetic tree *T* is a function that assigns a portion of the total edge length of *T* to each species. This can be seen as partitioning the total evolutionary history (the *phylogenetic diversity* [17]) of a phylogenetic tree among its species. Loosely speaking, these functions allocate the value of each edge length among that edge’s descendants in such a way that respects the symmetries of the tree shape. We write *φ*_*T, 𝓁*_ to denote the diversity index *φ* applied to the phylogenetic tree *T* given edge lengths by *𝓁*, although one or both of the subscripts may be omitted when clear from context. The formal definition of a diversity index builds on the related class of allocation functions [16]. Let *T* = (*V, E*) be a rooted phylogenetic *X*-tree with edge length assignment function *𝓁*. An *allocation function φ 𝓁* : *X* → ℝ^≥0^ is a real-valued function on the set of leaves of *T* that both satisfies the following equation:

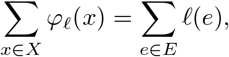

and moreover may be expressed as *φ 𝓁* (*x*) = Σ_*e*∈*E*_ *γ*(*x, e*) *𝓁* (*e*), | for every edge length assignment function *𝓁*, where all of the *coefficients γ*(*x, e*) are from the interval [0, 1]. Importantly, this means each such coefficient is non-negative.

A *diversity index* (on *T*) is an allocation function *φ* _*𝓁*_ : *X* → ℝ ^*>*0^ given by *φ 𝓁* (*x*) = Σ_*e*∈*E*_*γ*(*x, e*)*𝓁*(*e*) for every edge length assignment function *𝓁*, that additionally satisfies the conditions (DI_1_) and (DI_2_) below:

- (DI_1_) *Descent condition: γ*(*x, e*) = 0 if *x* is not descended from *e*.
- (DI_2_) *Neutrality condition:* The coefficients *γ*(*x, e*) are a function of the tree shape of *P*_*e*_. Moreover, suppose that *P*_*e*_ and *P*_*f*_ are pendant subtrees of *T* with the same tree shape. If the leaves *x* in *P*_*e*_ and *y* in *P*_*f*_ appear in corresponding positions in their respective subtrees, then *γ*(*x, e*) = *γ*(*y, f*).

One important consequence of these conditions is that *γ*(*x, e*) = 1 if *e* is the edge with leaf *x* as its terminal vertex. That is, the length of the edge incident to each leaf represents the evolutionary history unique to that leaf’s species, so its value must be allocated entirely to that species and no other.

We call *φ*(*x*) the diversity index *score* of *x* under *φ*. When comparing these scores we use, and repeat here, the definitions of *ranking, strict*, and *reversible* as they appear in [14] (pp. 2,3). A *ranking π*(*S, f*) for a set *S* = *{s*_1_, …, *s*_*n*_*}* based on a function *f* : *S* → ℝ is an ordered list of the elements of *S* such that *f* (*s*_*i*_) ≥ *f* (*s*_*j*_) if and only if *s*_*i*_ appears before *s*_*j*_ in *π*. A ranking *π*(*S, f*) is called *strict* if none of the values of *f* in *S* are equal. A ranking *π*_*T*_ is called *reversible* for diversity index *φ* if there is a subset *X* ^′^ of *X* whose removal from *T* leads to an induced subtree 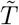 whose corresponding ranking 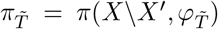 ranks the species in the opposite order to the ranking 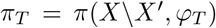. We call the removal of species from *X* an *extinction event* and will write 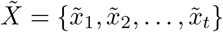 for the *t* species from *X* that survive an extinction event. The induced subtree 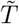 is the minimal subtree of *T* that spans 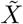 and the root vertex. Note that extinction events often cause vertices with out-degree one to appear. We suppress any such vertex and sum the lengths of its incident edges to give the length of the resultant edge in 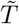.

It will be useful to discuss diversity indices on *T* in terms of their *ratios of allocations*, which we now define. Let 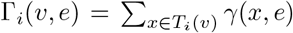, that is, the sum of all coefficients associated with both edge *e* and some leaf in the *i*-th pendant subtree below vertex *υ*. Let *e* = (*u, υ*) be an edge of *T*. Then the ratio Γ_1_(*υ, e*) : …: Γ_*d*_(*υ, e*) is called the *ratio of allocations* at *υ*, where *d* is the out-degree of *v*. Moreover, diversity indices in this paper will be assumed to be *consistent* in the sense that the values Γ_*i*_(*υ, f*) lie in the same ratio as the ratio of allocations at *υ* for every edge *f* in the path between the root vertex and *υ*. Each diversity index may be described by its (consistent) ratios of allocations, with [16] showing both how these ratios may be converted into *γ*(*x, e*)-type coefficients and that all diversity indices may be expressed in a consistent form.

Two types of diversity index will be considered further, defined in terms of their ratios of allocations. A *boundary* diversity index is a diversity index where it is possible for a term in a ratio of allocations to equal zero. In other words, boundary diversity indices contain ratios of allocations that mean some leaf is allocated no portion of the evolutionary history arising from one or more of its ancestral edges. We call any diversity index that is not boundary an *interior* diversity index. Equivalently, a diversity index *φ*(*x*) = Σ_*e*∈*E*_ *γ*(*x, e*)*𝓁*(*e*) is interior if and only if *γ*(*x, e*) is strictly positive whenever *x* is descended from *e*. The names ‘boundary’ and ‘interior’ refer to the position of a diversity index in *S*(*T, 𝓁*), the compact convex space of diversity indices on the tree *T* under edge length assignment *𝓁* (see [16] for details).

Given a rooted phylogenetic tree *T*, its non-root non-leaf vertices may be categorised by an equivalence relation ∼, where for *u, υ* ∈ *V* (*T*) we write *u* ∼ *υ* if and only if the multiset of tree shapes *{T*_*a*_(*u*), *T*_*b*_(*u*), …*}* is the same as the multiset of tree shapes *{T*_*a*_(*υ*), *T*_*b*_(*υ*), …*}*. Each diversity index can be thought of as a rule that determines ratios of allocations for every possible ∼-equivalence class. For a ∼-equivalence class with representative *υ*, we say that [*υ*] has *breadth* equal to |*c*_*T*_ (*υ*)|. Let a *singular* ∼-equivalence class be one whose representative vertex is the parent of a leaf of *T*. Equivalently, for a singular ∼-equivalence class with representative *υ*, the multiset of tree shapes *{T*_*a*_(*υ*), *T*_*b*_(*υ*), …*}* contains a tree that consists of a single vertex. In Figure 3, vertices *q, s* and *v* are from distinct singular ∼-equivalence classes. A *rigid diversity index* is a diversity index where the ratio of allocations is the same for at least two singular ∼-equivalence classes of different breadth on some rooted phylogenetic tree. We note that the majority of diversity indices are non-rigid interior indices. See Supplementary Material for details on the relative numbers of rigid, non-rigid and boundary indices.

### 2.1 Examples of diversity indices

In this section we present those diversity indices known from the literature and introduce some further examples to aid discussion. A small phylogenetic tree in Figure 3 is used to compare the effects of various diversity index functions. We begin with the formal definitions of the Fair Proportion (or Evolutionary Distinctiveness) index and the Equal-Splits index on a rooted phylogenetic *X*-tree *T*. For each leaf *x* ∈ *X*, the *Fair Proportion* (FP) index [2, 11] of *x* in *T* is given by

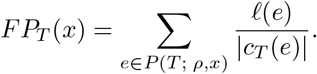

Now let *e* = (*u, υ*) be an edge of *T*. We define *π*(*e, x*) to be the product of the out-degrees of the interior vertices in the path *P* (*T* ; *υ, x*) unless *e* is a pendant edge, in which case *π*(*e, x*) = 1. Then for each leaf *x* ∈ *X*, the *Equal-Splits* (ES) index [1, 11] of *x* in *T* is given by

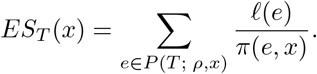

These two indices are both interior diversity indices, but are different in terms of being rigid or not, as shown in the following lemmas.

#### Lemma 1

*(i) The Equal-Splits diversity index is interior. (ii) The Fair Proportion diversity index is interior*.

*Proof* Let *υ* be an interior non-root vertex of a rooted phylogenetic *X*-tree *T*. Suppose *υ* has out-degree *d* and that *T*_1_(*υ*), *T*_2_(*υ*), …, *T*_*d*_(*υ*) are the maximal pendant subtrees descended from *υ*. Let |*X*| = *n* and write *n*_*i*_ for the number of leaves in *T*_*i*_(*υ*).

i. Under Equal-Splits the ratio of allocations at *υ* is 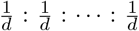. Each term in this ratio is non-zero for every choice of *υ*. Hence ES is an interior diversity index.
ii. Under Fair Proportion the ratio of allocations at *υ* is 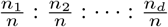. Each term in this ratio is non-zero for every choice of *υ*. Hence FP is an interior diversity index. □

#### Lemma 2

*(i) The Equal-Splits diversity index is rigid. (ii) The Fair Proportion diversity index is not rigid*.

*Proof* Let *T* be a rooted phylogenetic tree with at least two singular ∼-equivalence classes of different breadth. Let *u* and *υ* be vertices that are representatives of distinct ∼-equivalence classes where |*c*_*T*_ (*u*)| = *n*_*u*_ and |*c*_*T*_ (*υ*)| = *n*_*υ*_, and *n*_*u*_ *≠n*_*υ*_ .

i. Suppose *T* is binary. Then the ratio of allocations is 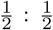 for every vertex under Equal-Splits. Thus the ratios of allocations at *u* and *υ;* are the same and hence ES is rigid.
ii. Under Fair Proportion the term in the ratio of allocations pertaining to a leaf that has parent *u* is 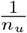. Similarly, a leaf with parent *υ* has the associated term of 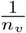 in the ratio of allocations at *υ*. As *n*_*u*_ *≠ n*_*v*_ these terms are different, and since these terms are necessarily the smallest terms in their respective ratios of allocations, the overall ratios must be different as well. Therefore FP is not rigid. □

Figure 2 pictorially represents the rigid/non-rigid classifications for ES and FP, as per Lemma 2. These two diversity indices are essentially the only ones found in existing biodiversity literature. We give a handful of further examples to outline the breadth of the diversity index concept and to illustrate some of the terms introduced above. The problem of finding useful diversity indices to complement FP and ES does not lie in finding functions that satisfy the diversity index definition, but rather in finding such functions that are biologically justified. The examples should therefore not all be interpreted as practical solutions to the problem of partitioning evolutionary history. For simplicity’s sake, we momentarily restrict our attention to rooted binary phylogenetic trees. Our examples are defined by a rule that determines their ratio of allocations at an arbitrary vertex *υ* with maximal descendant pendant subtrees *T*_*a*_(*υ*) and *T*_*b*_(*υ*). Table 1 displays the ratios of allocations when the example indices are applied to named vertices from the tree in Figure 3. Note that, by the neutrality condition of the diversity index definition, the vertices labelled *u, w* and *y* in Figure 3 will have a ratio of allocations of 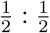 for every diversity index.

**Table 1:** Ratios of allocation for diversity indices discussed in Section 2.1, at labelled vertices from the tree *T* in Figure 3.

| Index | $p$ | $q$ | $r$ | $s$ | $t, v$ |
| --- | --- | --- | --- | --- | --- |
| FP | $\frac{5}{10} : \frac{5}{10}$ | $\frac{4}{5} : \frac{1}{5}$ | $\frac{3}{5} : \frac{2}{5}$ | $\frac{3}{4} : \frac{1}{4}$ | $\frac{2}{3} : \frac{1}{3}$ |
| ES | $\frac{1}{2} : \frac{1}{2}$ | $\frac{1}{2} : \frac{1}{2}$ | $\frac{1}{2} : \frac{1}{2}$ | $\frac{1}{2} : \frac{1}{2}$ | $\frac{1}{2} : \frac{1}{2}$ |
| $\alpha$ | $\frac{1}{2} : \frac{1}{2}$ | $\frac{2}{3} : \frac{1}{3}$ | $\frac{2}{3} : \frac{1}{3}$ | $\frac{2}{3} : \frac{1}{3}$ | $\frac{2}{3} : \frac{1}{3}$ |
| $\beta$ | $\frac{1}{2} : \frac{1}{2}$ | $1 : 0$ | $1 : 0$ | $1 : 0$ | $1 : 0$ |
| $\gamma$ | $\frac{1}{2} : \frac{1}{2}$ | $0 : 1$ | $0 : 1$ | $0 : 1$ | $0 : 1$ |
| $\delta$ | $\frac{1}{4} : \frac{3}{4}$ | $\frac{3}{4} : \frac{1}{4}$ | $\frac{1}{2} : \frac{1}{2}$ | $\frac{3}{4} : \frac{1}{4}$ | $\frac{3}{4} : \frac{1}{4}$ |
| $\epsilon$ | $\frac{3}{4} : \frac{1}{4}$ | $1 : 0$ | $1 : 0$ | $1 : 0$ | $\frac{1}{2} : \frac{1}{2}$ |
| $\zeta$ | $\frac{25}{50} : \frac{25}{50}$ | $\frac{16}{17} : \frac{1}{17}$ | $\frac{9}{13} : \frac{4}{13}$ | $\frac{9}{10} : \frac{1}{10}$ | $\frac{4}{5} : \frac{1}{5}$ |
| $\eta$ | $\frac{1}{2} : \frac{1}{2}$ | $\frac{1}{2} : \frac{1}{2}$ | $\frac{1}{2} : \frac{1}{2}$ | $\frac{3}{4} : \frac{1}{4}$ | $\frac{2}{3} : \frac{1}{3}$ |
| $\theta$ | $\frac{4}{8} : \frac{4}{8}$ | $\frac{8}{8} : \frac{0}{8}$ | $\frac{5}{8} : \frac{3}{8}$ | $\frac{0}{8} : \frac{8}{8}$ | $\frac{2}{8} : \frac{6}{8}$ |

**Fig. 2:**
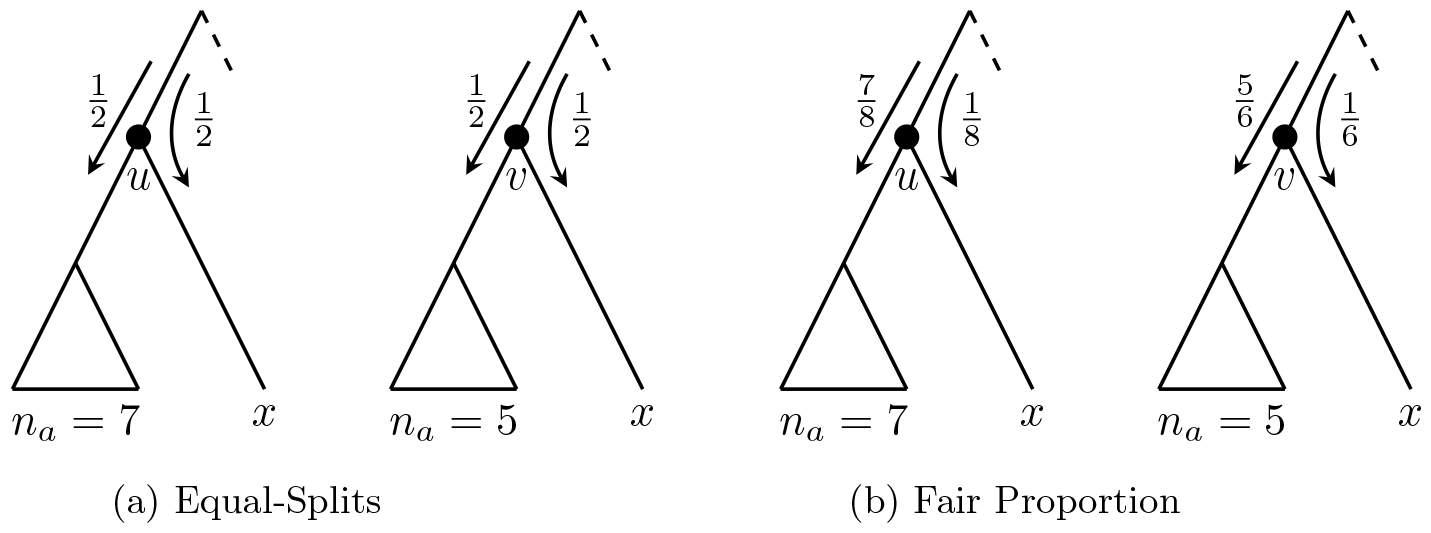
(a) The Equal-Splits index is a rigid diversity index, as the ratio of allocations is unchanged from *u* to *υ* although the number of leaves in the left-hand subtree (represented by a triangle) is changed. (b) In contrast, the Fair Proportion index is not a rigid diversity index, as can be seen from the different ratio of allocations in each case.

**Fig. 3:**
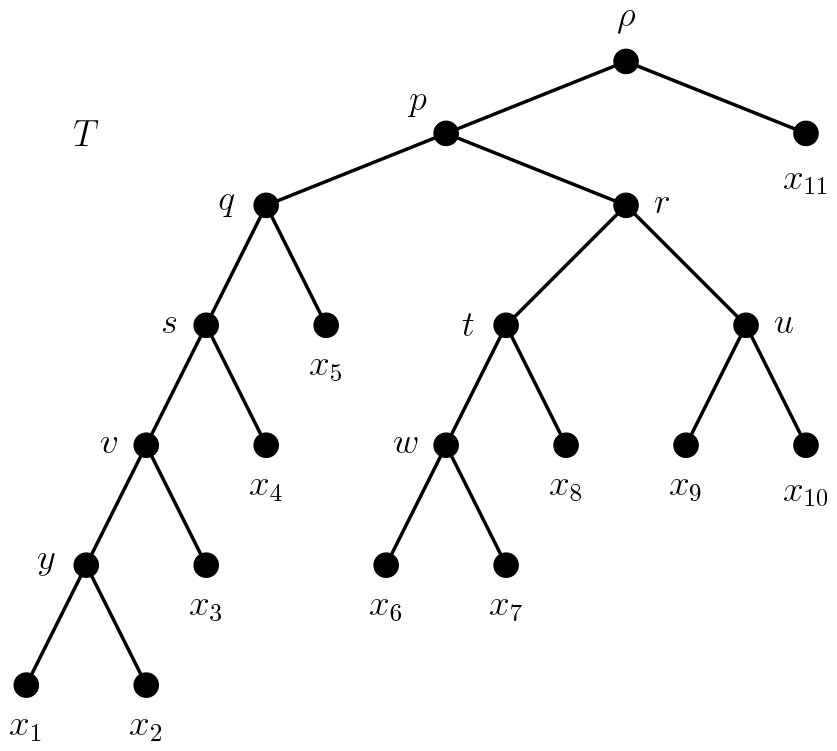
A rooted binary phylogenetic tree *T*. Many diversity indices may be defined on *T* by specifying ratios of allocations at vertices *p, q, r, s, t* and *υ*.

Our first index, *α*, compares the two pendant subtrees *T*_*a*_(*υ*) and *T*_*b*_(*υ*) and prioritises the subtree that contains more leaves. If both *T*_*a*_(*υ*) and *T*_*b*_(*υ*) contain the same number of leaves, the *α* index has a 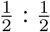 ratio of allocations at *υ*. Otherwise *α* has a 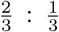 ratio of allocations at *υ* that allocates 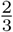 the preceding evolutionary history to the more populous subtree. Our second index, *β*, follows the same approach, but takes the ratios to an extreme. Again, if both *T*_*a*_(*υ*) and *T*_*b*_(*υ*) contain the same number of leaves, the *β* index has a 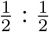 ratio of allocations at *υ*. Otherwise *β* has a 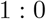 ratio of allocations at *υ* that allocates all of the preceding edge’s length to the more populous subtree. Another similar diversity index, *γ*, can be constructed by taking the other extreme, that is where each 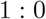 ratio of allocations gives all of the preceding edge’s length to the less populous subtree. The indices *α, β* and *γ* can be seen as three functions from a family of diversity indices that includes ES as its midpoint. All are rigid diversity indices, as the same ratio is used for many ∼-equivalence classes. Indices *β* and *γ* are also boundary diversity indices.

We can also choose other aspects of a tree’s structure on which to base a diversity index’s ratios of allocations. For example, the index *δ* uses the number of cherries rather than the number of leaves: if both *T*_*a*_(*υ*) and *T*_*b*_(*υ*) contain the same number of cherries, the *δ* index has a 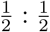 ratio of allocations at *υ*. Otherwise *δ* has a 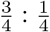 ratio of allocations at *υ* that allocates 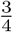 to the subtree with the greater number of cherries. (The value of 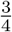 was chosen arbitrarily from the range [0, 1].) A further feature that could be used is the number of interior vertices in *T*_*a*_(*υ*) and *T*_*b*_(*υ*) that are parents to exactly one leaf. This feature is used by the *E* index, where the ratios of allocations are proportionate to the number of such vertices in each maximal pendant subtree descended from *υ*, or a 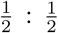 ratio if no such vertices exist. The use of proportionate allocations in this way is closer to how the Fair Proportion index works than the use of fixed ratios in *α, β, γ* and *δ*.

In addition to direct proportions of leaves, cherries, interior vertices or other features of tree topology, we can form ratios of allocations based on some function of these numbers. The (non-rigid) *ζ* diversity index uses the proportions of the squares of the numbers of leaves in each subtree to determine its ratios of allocations. We can also combine existing diversity indices piecewise. For example, our *η* index has ratios of allocations that match ES for vertices that have five or more descendant leaves and ratios that match FP for each other vertex. A final type of diversity index that we mention here is the arbitrary diversity index *θ*. We sampled five integers between 0 and 8 and used each as the numerator of the left hand term in the ratios in our table. This is clearly not a biologically-relevant approach but does determine a legitimate diversity index. Lastly we note that linear combinations of diversity indices are themselves diversity indices, subject to some basic constraints [16].

The indices above highlight the wide variety of possibilities using the diversity index concept. Each diversity index carries with it a set of assumptions about how evolutionary history is shared or embodied among descendants. The usefulness of each index depends on how biologically credible these assumptions are, because the differences in assumptions are reflected in the scores given to each species. For example, the indices in Table 1 give rise to the following scores for leaf *x*_3_ if all edges have unit length: *FP* (*x*_3_) = 1.88, *ES*(*x*_3_) = 1.94, *α*(*x*_3_) = 1.78, *β*(*x*_3_) = 1, *γ*(*x*_3_) = 2, *δ*(*x*_3_) = 1.61, *E*(*x*_3_) = 2.88, *ζ*(*x*_3_) = 1.63, *η*(*x*_3_) = 1.77 and *θ*(*x*_3_) = 1.75 (all rounded to 2 d.p.). Note the large range of values, from 1 to 2.88 units.

The Fair Proportion index assumes that each species descended from an edge exhibits that edge’s evolutionary developments equally. For Equal-Splits the assumptions are that at the time of the speciation event that terminated an edge, that edge’s developments were embodied equally among the new lineages, and that further speciation events do not alter this original separation. Index *β* effectively considers the less populous subtrees to be developing evolutionary history from scratch after a speciation event, whereas *γ* places this assumption on the more populous subtrees. We suggest that while the assumptions for FP and ES are reasonable, the final two are not readily justified.

It turns out that the *β* index can be more robust than interior indices (see Section 4, page 18 for details). Balanced against this observation is the fact that often it achieves this robustness by essentially ignoring much of the tree structure. Many *β* index scores are simply pendant edge lengths. This circumvents the entire reason for using phylogenetic trees to give a more structured and nuanced picture of evolution compared to plain distance methods. Hence we do not see *β* as a comprehensive solution to the robustness problem. Boundary indices can also act in an inconsistent and somewhat arbitrary fashion. A zero term in a ratio of allocations is making a strong claim about the way evolutionary history is embodied in living species. Each zero allocation excludes some species from a share of ancestral evolutionary developments, despite these species being descended from the ancestor where such developments arose. We therefore conclude that boundary diversity indices are not a good model for measuring the evolutionary history of species and are only interesting as the theoretical limit of the diversity index concept.

## 3 Non-rigid interior diversity indices

We begin our examination of robustness with a broad question: Given the freedom to choose any positive edge lengths for a known rooted phylogenetic tree, is there a set of extinctions that induces a strict and reversible ranking on the surviving species’ diversity index scores? Note that eventually, after more and more extinctions, we must arrive at a set of surviving leaves that form a strict and reversible ranking, regardless of the particular diversity index (even if this is a trivial set containing one or two leaves). Hence the pertinent question is: How many extinctions are necessary and sufficient to achieve this effect? The answer to this question depends on the diversity index being used and the structure of the given phylogenetic tree. For the FP index on rooted binary phylogenetic trees the extinction of one leaf per cherry is both necessary and sufficient [14, Theorem 2]. In this section we show that all non-rigid interior diversity indices exhibit the same ranking reversal behaviour as FP. That is, the number of necessary and sufficient extinctions, and their distribution among the leaves, is the same for any non-rigid interior index as it is for FP. Our results are given for rooted phylogenetic trees with unrestricted out-degree.

The separation of diversity indices into the classes of rigid and non-rigid is needed because the patterns of extinctions necessary to generate a diversity index ranking reversal differ in each case. The number of necessary and sufficient extinctions for rigid interior diversity indices is somewhat larger, and includes those extinctions necessary in the non-rigid case. Hence, rigid interior indices are slightly more robust than non-rigid indices, although still susceptible to complete ranking reversals as shown in Section 4.

The first result we establish concerns the number and type of extinctions necessary for a non-rigid index ranking reversal. The question of sufficiency will then be addressed, showing that the necessary extinctions suffice. Theorem 1 of [14] described the necessary number and type of leaf deletions for the Fair Proportion index on rooted binary phylogenetic trees. In fact, their result that at least one leaf per cherry must be deleted generalises directly to every diversity index. We now present a generalisation of that theorem, covering all phylogenetic diversity indices and also those rooted phylogenetic trees that are not binary. Proposition 3 includes the one-per-cherry extinction events as a special case and is proven in a very similar manner to Theorem 1 of [14].

### Proposition 3

*Let T be a rooted phylogenetic X-tree with* |*X*| ≥ 3 *and let φ be a diversity index on T*. *Suppose π*_*T*_ *is a strict and reversible ranking concerning the φ diversity index with respect to* 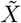 *and induced subtree* 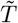. *Then* 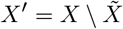 *contains at least all but one of the leaves adjacent to each interior vertex of T* .

*Proof* With a view to contradiction, assume that leaves *x*_*i*_ and *x*_*j*_ are adjacent to the interior vertex *v* and that neither *x*_*i*_ nor *x*_*j*_ is in *X* ^*′*^. Let *e*_*i*_ and *e*_*j*_ be the pendant edges incident with *x*_*i*_ and *x*_*j*_ respectively. By the neutrality condition in the definition of a diversity index, *φ*_*T*_ (*x*_*i*_) = *p*+*𝓁*(*e*_*i*_) and *φ*_*T*_ (*x*_*j*_) = *p*+*𝓁*(*e*_*j*_) for some real value *p*, and 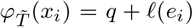 and 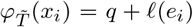 for some real value *q*. Without loss of generality suppose that *π*_*T*_ (*x*_*i*_) *< π*_*T*_ (*x*_*j*_) and 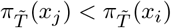. But then substituting the values from above gives rise to the contradictory expressions *𝓁*(*x*_*i*_) *< 𝓁*(*x*_*j*_) and *𝓁*(*x*_*j*_) *< 𝓁*(*x*_*i*_). Hence our initial assumption was false and thus at least one of *x*_*i*_ and *x*_*j*_ is in the set *X*^*′*^. As *x*_*i*_ and *x*_*j*_ were chosen arbitrarily from the leaves adjacent to *υ*, the result follows. □

Figure 4 shows a rooted phylogenetic tree on nine leaves, where the leaves are grouped according to their parent vertex. The unfilled leaves represent one set of leaf deletions of the smallest size required for a strict and reversible ranking to be possible. This number of deletions can be a large proportion of the leaves for a nonbinary tree. However, it can be much smaller for some rooted binary phylogenetic trees, such as a single vertex for caterpillar trees.

**Fig. 4:**
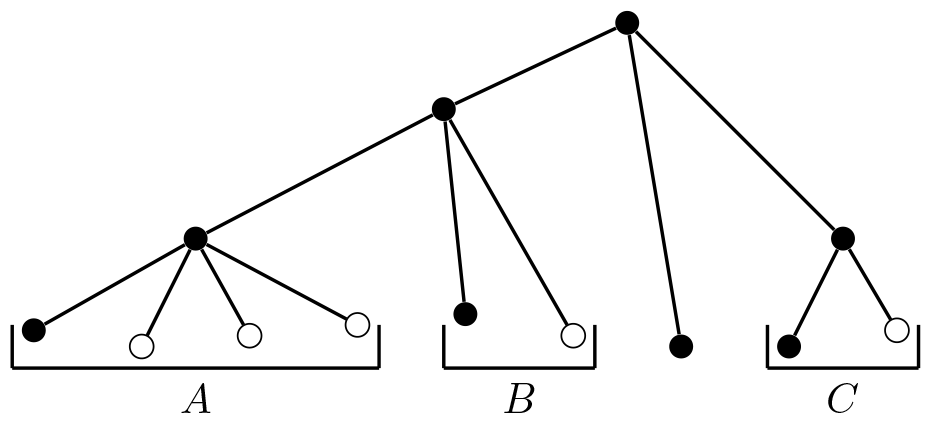
At least three out of the four leaves in *A* and one of the two leaves in each of *B* and *C* must be deleted in order for a strict reversal of diversity index score values to be possible. One such set of leaves is indicated above by the unfilled vertices.

Our task now is to show that this necessary set of extinctions is also sufficient to cause a non-rigid index ranking reversal. This was proven for the Fair Proportion index, as Theorem 2 in [14]. The main result of this section is to show that that theorem can be extended beyond the FP index to apply to every non-rigid interior diversity index (Theorem 5). Hence, FP is not uniquely subject to robustness issues in light of ongoing extinctions.

Let *φ* be an interior non-rigid diversity index. Our inductive proof below relies on building up a collection of edge lengths for a phylogenetic tree *T* based upon edge lengths that make *φ* reversible on the maximal pendant subtrees of *T*. This follows the same general approach of Fischer and colleagues in the proof of their Theorem 2 [14]. Both results are proved inductively, building from small trees into larger ones. As the smaller trees are combined into larger ones and eventually into *T* as a whole, the edge lengths are repeatedly adjusted to have strict and reversible rankings at each step. The main difference here is that our approach abstracts the scores away from any particular diversity index and tree structure that creates them. We only require that all, or all but one, species experience a change in diversity score because of an extinction event. We begin with Lemma 4, a result that allows us to adjust edge lengths from the smaller trees in a useful way throughout the proof of Theorem 5.

### Lemma 4

*Let T* = (*T*_*a*_, *T*_*b*_, …) *be a rooted phylogenetic X-tree where* |*X*_*a*_| ≥ 2. *Let φ*_*T*_ = Σ_*e*∈*E*(*T*)_ *γT* ^(*x, e*)*𝓁*(*e*)^ *be an interior diversity index on T that is not rigid. Suppose that φ induces a strict and reversible ordering of taxa in* 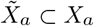 *after the extinction of all but one of the leaves adjacent to each interior vertex of T*_*a*_. *Let* 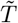 *be the rooted phylogenetic subtree of T induced by this extinction event. Then there exist edge lengths for T such that* 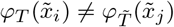 *for all* 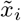 *and* 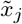 *in* 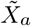.

*Proof* Let 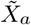 be indexed in such a way to give the *φ* diversity index score ranking 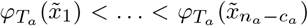, where *c*_*a*_ is the number of leaves deleted under this extinction event. We prove the result by describing how those edge lengths of *T*_*a*_ that allow the ranking reversal on 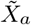 can be adjusted to give distinct *φ* index scores.

First, assume that 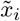 is a leaf with 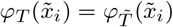. As *φ* induced a strict and reversible ranking, 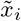 is the only leaf for which this equality holds. Let *e*_*i*_ be the pendant edge incident to 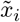 in *T* and let 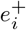 be the edge in *T* whose terminal vertex is the parent of 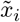. Suppose that 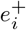 is edge *a*, incident with the root of *T*. Since *φ* is an interior diversity index, 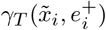 and 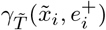 are both strictly positive. Moreover, as *φ* is not rigid we have 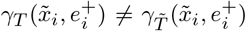 and because *x*_*i*_ is the only descendant of *e*_*i*_, we have 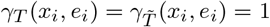. But then

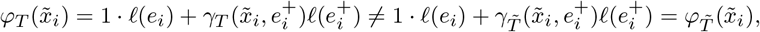

contradicting our supposition.

So it must be the case that 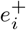 lies in *T*_*a*_. Now extending 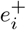 will increase 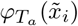 and 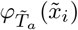 by different amounts because *φ* is not rigid. Specifically, we choose to extend 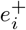 by the lesser of 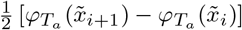 and 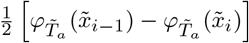 (or the defined value from these two choices if only one exists). This particular choice is small enough to ensure that neither the *π*_*a*_ ranking nor the 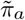 ranking is altered by the extension. Hence strict reversibility is maintained. Moreover, extending 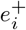 in such a manner ensures that 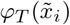 and 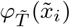 will have distinct values.

Second, assume that 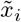 and 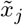 are distinct leaves with 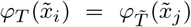. Then we can extend *e*_*i*_, the pendant edge incident to 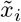, by the lesser of 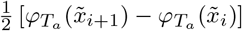 and 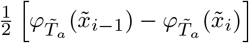 (or the defined value from these two choices if only one exists). Extending the pendant edge incident to 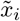 increases both 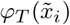 and 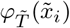 by equal amounts but does not alter 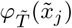. This particular choice is small enough to ensure that neither the *π*_*a*_ ranking nor the 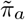 ranking is altered by the extension. Hence strict reversibility is maintained. Moreover, extending the pendant edge incident to 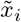 in such a manner ensures that 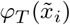 and 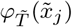 will have distinct values. We repeat this step as necessary for every such pair of leaves.

Finally, we choose *𝓁*(*a*) to be short enough to ensure that it is less than the smallest difference between any two 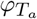 or 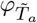 index scores from our modified version of *T*_*a*_. So even if one leaf were to be allocated the entire length of edge *a*, the ranking orders *π*_*a*_ and 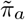 would not change.

Therefore, given a set of edge lengths for *T*, we can adjust them in the manner described above to ensure 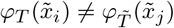 for all 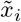 and 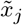 in 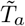. □

We turn now to the first main result of the paper.

### Theorem 5

*Let T* = (*T*_*a*_, *T*_*b*_, …) *be a rooted phylogenetic X-tree with* |*X*| = *n, where n* ≥ 2. *Let* 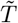 *be the induced* 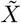*-tree that results from deleting all but one of the leaves adjacent to each interior vertex of T*. *Then, for any non-rigid interior diversity index φ, there exists an edge length assignment function 𝓁 on T such that there is a strict φ ranking π*_*T*_ *for the leaves of T that is reversible with respect to* 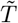 *and such that* 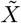 *contains the species that has the highest φ index score in T* .

*Proof* We first prove the result for a rooted phylogenetic tree where the root vertex has out-degree 2 before describing how this technique can be extended to trees with larger out-degree at the root. We proceed by induction on the number of interior vertices present in a rooted phylogenetic tree. As a base case we consider a rooted phylogenetic ‘star’ tree with exactly one interior vertex, the root, and *n* edges of different lengths. If every taxon except that incident with the longest edge is deleted, then this remaining taxon is in a strict and reversible ordering (of size one) and the result holds.

Next, for the inductive step assume that the theorem holds for the non-rigid interior diversity index *φ* on all rooted phylogenetic trees with less than *k* interior vertices. Let *T* = (*T*_*a*_, *T*_*b*_) be a rooted phylogenetic tree with *k* interior vertices and *n* leaves, as drawn in Figure 1. Without loss of generality we have *n*_*a*_ ≥ *n*_*b*_. Let *c*_*T*_ be the number of leaves removed when deleting all but one of the leaves adjacent to each interior vertex of *T* and let *c*_*a*_ and *c*_*b*_ denote the number of these leaves contained in *T*_*a*_ and *T*_*b*_, respectively. We show that by using the edge lengths that allowed reversible orderings of *φ* on *T*_*a*_ and *T*_*b*_, we are able to choose edge lengths that make *φ* reversible on *T* as a whole.

As *T*_*a*_ has less than *k* interior vertices, the extinction of all but one of the leaves adjacent to each interior vertex of *T* induces a strict and reversible ranking *π*_*a*_ on the leaves of *T*_*a*_. Let 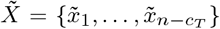 be the set of leaves that is not deleted, labelled such that 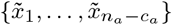 is contained in *T*_*a*_ and 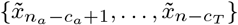 is contained in *T*_*b*_. Relabelling if necessary, we have

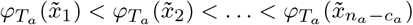

and

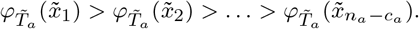

As described in Lemma 4, we choose a small value for *𝓁*(*a*) that does not alter any of these inequalities. Then we construct a set of combined *φ* index scores for 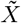 from both *T* and 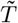, denoted Σ_*a*_ ⊂ ℝ^*>*0^. In particular:

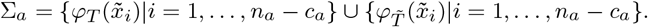

By Lemma 4, since *φ* is non-rigid and interior, each element of Σ_*a*_ can be made to have a unique value by adjusting edge lengths within *T*_*a*_. Moreover, this is achievable in a way that the ranking *π*_*T*_ restricted to *T*_*a*_ is the same as *π*_*a*_. Thus we are able to ensure that Σ_*a*_ contains 2(*n*_*a*_ − *c*_*a*_) distinct positive real numbers without altering the ranking that we began with. See Figure 5 for an illustration of this construction. Let Σ_∗_ be the subset of Σ_*a*_ containing the *n*_*a*_ − *c*_*a*_ least elements of Σ_*a*_ under the usual ordering of real numbers and let Σ^∗^ be the subset of Σ_*a*_ containing the *n*_*a*_ − *c*_*a*_ greatest elements of Σ_*a*_ under the same ordering. Hence, Σ_*a*_ is the disjoint union of Σ_∗_ and Σ^∗^.

**Fig. 5:**
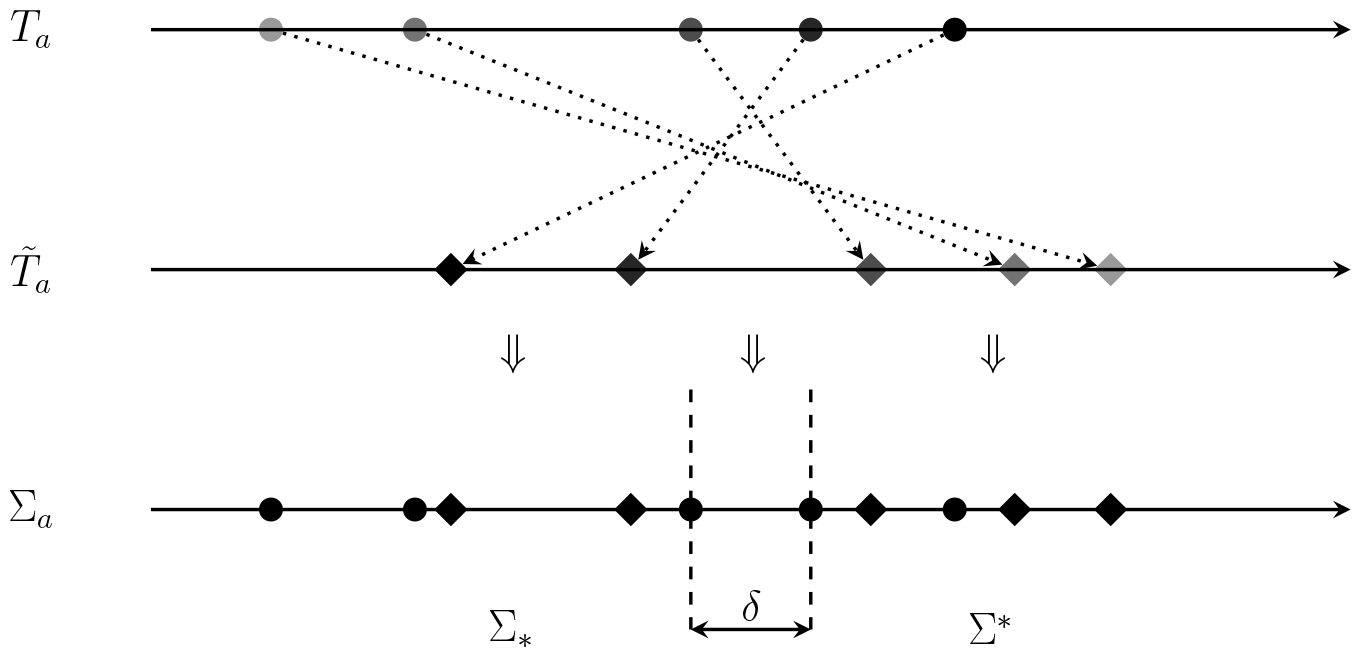
Illustration of the first part of the construction described in the proof of Theorem 5. Each horizontal line represents the set of real numbers. Dotted lines indicate the reversal of index scores of leaves in *T*_*a*_ after the extinction of all but one of the leaves adjacent to each interior vertex of *T*, as per our induction hypothesis. The scores for both *T*_*a*_ and 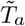 are combined on the bottom number line, where, by Lemma 4, the 2(*n*_*a*_ − *c*_*a*_) scores may be made distinct without affecting the initial reversal. Finally the edge lengths of subtree *T*_*b*_ are adjusted and scaled to fit the index scores into the interval of size *δ* between the vertical dashed lines.

Next we consider the remaining taxa of *T*. The subtree *T*_*b*_ has less than *k* interior vertices and hence the extinction of all but one of the leaves adjacent to each interior vertex of *T* induces a strict and reversible ranking *π*_*b*_ on 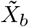. Similar to *T*_*a*_, relabelling if necessary, we have

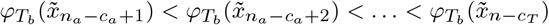

and

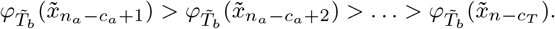

As in the *T*_*a*_ case, we choose a small enough value for *𝓁*(*b*) so that the above rankings are maintained. We construct the set

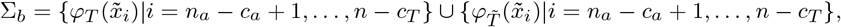

but, in contrast to Σ_*a*_, do not require that Σ_*b*_ contains 2(*n*_*b*_ −*c*_*b*_) distinct values. (For instance *T*_*b*_ may consist of a single vertex whose diversity index score is necessarily fixed as the length of its pendant edge.) However, we do note that, because diversity indices allocate the entirety of each pendant edge length to their incident leaf, every member of Σ_*b*_ is strictly positive.

Now let *δ* = min Σ^∗^ − max Σ_∗_ be the size of the gap between the two halves of Σ_*a*_ (see Figure 5). We uniformly multiply the lengths of edges in *E*(*T*_*b*_) ∪ *{b}* by a positive real constant *c*, chosen so that *c* (max Σ_*b*_) *< δ*. That is, we scale (down) the entirety of *T*_*b*_ as well as edge *b* so that the range of values in the rescaled Σ_*b*_ extends less than *δ*. (In practice it may be tidier to scale up *T*_*a*_ and *a* to achieve the same result.) Call the resulting subtree 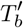. Finally, to each pendant edge in 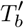 add the value of max Σ_∗_. This shifts the index scores of 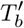 into the gap between Σ_∗_ and Σ^∗^ without altering the ranking of these leaves.

Let 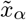 be the taxon such that 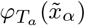 is the largest value in Σ_∗_ among scores from *T*_*a*_ (as opposed to 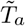). The result of the above construction is that we have

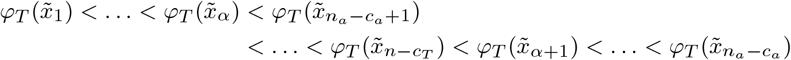

and

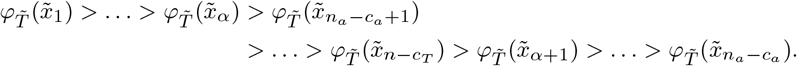

Therefore the extinction of 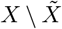 has induced a strict and reversible ranking on 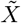 given our chosen edge lengths. Hence the induction is proved and the theorem holds for all rooted phylogenetic trees where the root has out-degree 2.

We now describe how to extend the above idea to phylogenetic trees where the root has greater out-degree. Let 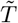 be the rooted phylogenetic tree induced from *T* by the extinction of all but one leaf adjacent to each interior vertex. Suppose that 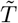 has maximal pendant subtrees 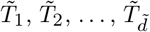, where 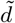 is the out-degree of the root in 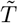. Without loss of generality we further suppose that 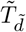 is the maximal pendant subtree with the fewest leaves from this list. Note that, since 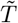 contains at most one leaf adjacent to each interior vertex, at most 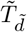 consists of a single leaf.

We first apply the above interleaving process to the subtrees 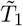 and 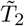, with one small change. Since 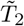 has at least two leaves we use Lemma 4 to ensure that the set Σ_2_ contains no repeated values. Thus the strict and reversible ranking obtained across leaves from 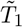 and 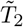 contains some interval *δ* ^*′*^ between the lower half of diversity index scores and the upper half of these scores. It is into this interval, scaling as before, that we place the scores from Σ_3_, which itself will contain an inter-score interval *δ* ^*″*^ in which to place scores from Σ_4_, and so on. We proceed iteratively in this manner until finally we place the scores from 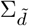 into the last interval. The result is a single completely reversible ranking across all leaves of 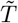, with edge lengths chosen according to the steps described. Therefore the statement of the theorem is proved for all rooted phylogenetic trees. □

## 4 Rigid interior diversity indices

Theorem 5 has shown that the non-rigid interior diversity indices, to the same extent as FP, are not robust to extinctions. We would also like to establish the same understanding of rigid interior diversity indices. That is, which extinctions are required before a complete ranking reversal can occur and, given the freedom to choose positive edge lengths, which extinctions are sufficient? For a ranking derived from a rigid interior diversity index to undergo a strict reversal, the extinction of at least all but one leaf adjacent to each interior vertex is still required, as Proposition 3 includes all diversity indices. However, as an examination of (the quintessential rigid diversity index) ES will show, such an extinction event may not be sufficient to reverse the entire ranking of survivors’ index scores (Proposition 6). We then establish the necessary and sufficient sets of extinctions for this class of diversity index. This involves a new definition, categorising some leaves of a rooted phylogenetic tree as ‘isolated’. We show that the additional extinction of these isolated leaves is sufficient for inducing a reversible index ranking for any rigid diversity index (Theorem 9).

### Proposition 6

*Let T be a rooted caterpillar tree with at least four leaves. Then the extinction of one leaf from the cherry of T is necessary, but not sufficient, to cause a strict ranking of Equal-Splits index scores to reverse*.

*Moreover, let X* = *{x*_1_, *x*_2_, *x*_3_, …, *x*_*n*_*} be the leaves of T, where x*_*n*_ *is adjacent to the root vertex, and there is a path of length two from the root vertex to x*_*n*−1_. *Then the extinction of any proper subset of {x*_1_, *x*_2_, *x*_3_, …, *x*_*n*−2_*} cannot cause a strict ranking of Equal-Splits index scores to reverse*.

*Proof* Let *x*_1_ and *x*_2_ be the two distinct leaves that form the unique cherry of *T*. By Theorem 1 in [14], it is necessary for at least one of *x*_1_, *x*_2_ to be deleted for the ranking of ES index scores to reverse.

The extinction of, say, *x*_1_ alone will lead to an increase in the ES index score of *x*_2_, but the ES index scores of all other vertices remain the same. If *T* contains at least four leaves, there are at least two leaves whose index scores are unaffected, and hence their ranking order does not reverse. Thus the extinction of *x*_1_ or *x*_2_ alone is not enough to reverse the ES index scores.

For a rooted caterpillar tree with *n* ≥ 4 leaves, suppose a strict subset of *{x*_1_, *x*_2_, *x*_3_, …, *x*_*n*−2_*}* becomes extinct. Note that the extinction of a leaf *x*_*i*_ affects the ES index score of precisely those leaves descended from the parent of *x*_*i*_ and no others. By this reasoning the ES index scores of *x*_*n*−1_ and *x*_*n*_ are unaffected by such an extinction and the ranking cannot reverse. Therefore the deletion of *x*_*n*−1_ or *x*_*n*_ is necessary for an ES ranking reversal. □

This result confirms our claim above, that some rigid interior indices can be robust to the type of extinction events that cause ranking reversals on nonrigid interior indices. We would like to know how few extinctions beyond those specified in Proposition 3 are necessary to induce a ranking reversal. Let *T* be a phylogenetic *X*-tree rooted at *ρ* and let 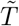 be the 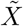-tree induced by the extinction of 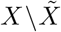. A *fixed leaf* for the diversity index *φ* on *T* is any leaf *x* in *X* where 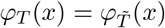 independent of the edge length assignment. For a strict and reversible *φ* ranking on 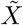 to exist, there must be at most one fixed leaf for *φ* on *T*. The focus on fixed leaves can help us to understand the robustness of various diversity indices. Consider the *β* diversity index defined in Section 2.1. When applied to caterpillar trees with *n* leaves (see *Cat*_*n*_ in Figure 6), we require more than *n* − 4 extinctions to possibly induce a *β* ranking reversal on surviving leaves. This is because all leaves on *Cat*_*n*_, except the three leaves furthest from the root, are fixed leaves under *β* for every extinction event that they survive. Thus it is necessary for at least *n* − 4 of these fixed leaves to go extinct, plus one from the cherry. This makes *β* more robust on *Cat*_*n*_ than other indices we examine, however this advantage needs to be weighed against the disadvantages of boundary indices discussed earlier.

**Fig. 6:**
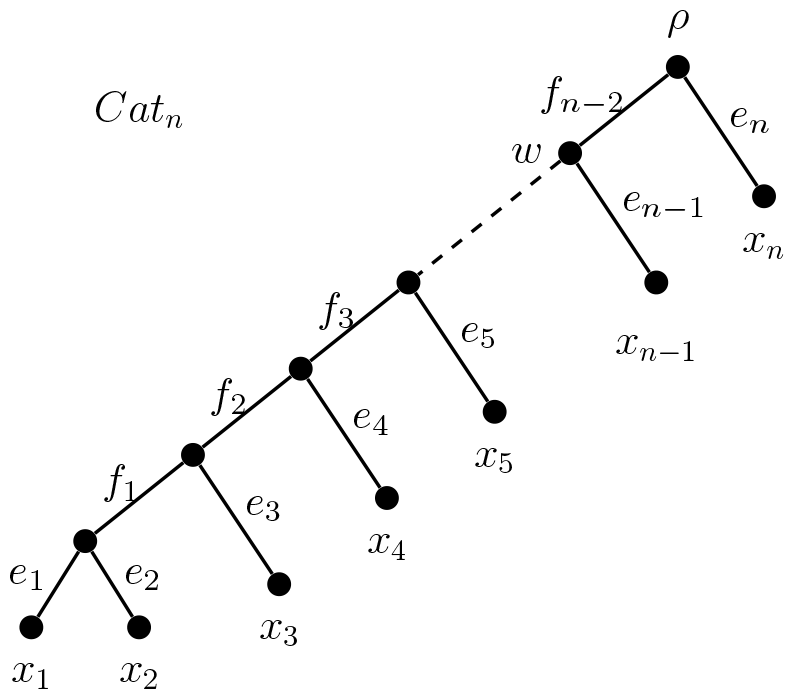
A general caterpillar tree used to show that ES index score rankings can be reversed with as few as two extinctions. Observe that leaf *x*_*n*−1_ is the only isolated leaf in this tree.

Next, let *x* ∈ *X* be a leaf vertex with parent *υ* ∈ *V* (*T*) distinct from *ρ*, where *x* is not contained in any *m*-cherry. Then we call *x* an *isolated leaf* if there is no vertex *u* in the path *P* (*T* ; *ρ, υ*) \ *{ρ, υ }*, such that *u* itself is the parent of a leaf vertex. For example, in the tree *Cat*_*n*_ in Figure 6, leaf *x*_*n*−1_ is an isolated leaf because there is no additional vertex in the path from the root *ρ* to *w*, the parent of *x*_*n*−1_, other than the endpoints. Leaf *x*_*n*_ is not isolated because its parent vertex is the root. All other leaves of this tree are not isolated because the vertex *w* appears on all paths from *ρ* to the parents of other leaves. Proposition 7 below shows that in addition to all but one leaf adjacent to each interior vertex, we also require all but one isolated leaf to go extinct before a strict and reversible ranking on *T* can occur for rigid interior diversity indices.

### Proposition 7

*Let T be a rooted phylogenetic X-tree with* |*X*| ≥ 3 *and let φ be a rigid diversity index on T*. *Suppose π*_*T*_ *is a strict and reversible ranking concerning the φ diversity index for T with respect to* 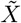 *and induced subtree* 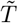. *Then* 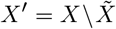 *contains at least all but one of the leaves adjacent to each interior vertex of T and at least all but one of the isolated leaves of T*. *In the case that one of the maximal pendant subtrees of T contains a single leaf, then X* ^*′*^ *must contain all of the isolated leaves of T* .

*Proof* Each isolated leaf may be a fixed leaf if *φ* is a rigid diversity index. Any leaf whose parent vertex is the root is also a fixed leaf, because its diversity index score is always just the length of the incident pendant edge. To induce a strict and reversible ranking on 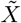 under a diversity index at most one fixed leaf for *φ* on *T* can remain. Therefore, the deletion of at least all but one isolated vertex is required and if one of the maximal pendant subtrees of *T* contains a single leaf then the final isolated leaf must be deleted too. Since the extinction of at least all but one of the leaves adjacent to each interior vertex is needed to reverse any strict diversity index ranking, the result is shown. □

Our next results show that the extra extinction of isolated vertices is sufficient to give a ranking reversal for rigid interior diversity indices. That is, those interior diversity indices not covered by Theorem 5 are also not robust given this larger set of extinctions. Lemma 8 describes how we can choose edge lengths for a rooted phylogenetic tree *T* so that, after the extinction of all isolated leaves and all but one leaf adjacent to each interior vertex of *T*, we can ensure that there is at most one fixed leaf for any rigid interior diversity index. Theorem 9 then uses this lack of multiple fixed leaves to determine edge lengths that induce a ranking reversal.

### Lemma 8

*Let T be a rooted phylogenetic X-tree and let φT* = Σ_*e*∈*E*(*T*)_ *γ*(*x, e*)*𝓁*(*e*) *be an interior diversity index on T*. *Suppose that* 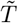 *is the rooted phylogenetic* 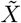*-tree induced by the extinction of all isolated leaves and all but one leaf adjacent to each interior vertex of T*. *Then there exist edge lengths for T such that* 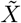 *contains at most one fixed leaf for φ on T* .

*Proof* We prove the result by describing how certain edge lengths can be extended to ensure that at most one fixed leaf for *φ* exists in *T*. Let *𝓁* be an edge length assignment function on *E*(*T*). Assume that there exists vertex 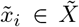 such that 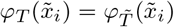 and 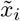 is not adjacent to the root vertex. If 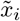 is a member of an *m*-cherry in *T*, let 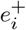 be the edge whose terminal vertex is the parent of 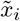 in *T*. Then increasing the length of 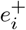 will increase 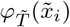 by *m* times the amount that 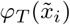 increases. Hence, 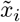 is no longer a fixed leaf after extending 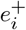.

If 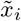 is not a member of an *m*-cherry in *T*, then there exists some isolated leaf *x*_*j*_ ∈ *X* whose parent in *T* is in the path 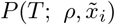. This must be the case or 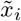 would itself be an isolated leaf of *T*, contradicting the fact that all isolated leaves have been deleted. Let 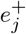 be the edge in *T* whose terminal vertex is the parent of *x*_*j*_. We set 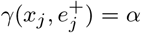, where *α* lies in the open interval (0, 1) because *φ* is not a boundary index. Let 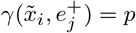. Since *φ* has a consistent form (Proposition 11 of [16]), we can write *p* = (1 − *α*)*q* for some *q* ∈ (0, 1). Increasing the length of 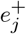 by, say, *k* units increases 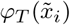 by *pk* but increases 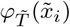 by *qk*, a different amount since *α* is nonzero. Hence 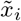 is no longer a fixed leaf after extending 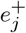.

Therefore we can extend particular edges as required to get a set of edge lengths for which 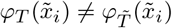 whenever 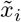 is not adjacent to the root vertex. □

### Theorem 9

*Let T be a rooted phylogenetic X-tree and let φ*_*T*_ = Σ_*e*∈*E*(*T*)_ *γ*(*x, e*)*𝓁*(*e*) *be an interior diversity index on T*. *Suppose that* 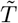 *is the rooted phylogenetic* 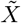*-tree induced by the extinction of all isolated leaves and all but one leaf adjacent to each interior vertex of T*. *Then there exists an edge length assignment function 𝓁 on T such that there is a strict φ ranking π*_*T*_ *for the leaves of T that is reversible with respect to* 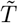.

*Proof* The proof consists of two parts. We first construct an edge length assignment function *𝓁* ^*′*^ for *T*. Next, we adjust the lengths of pendant edges to construct the edge length assignment function *𝓁* with the desired reversal property.

Let *F*_iso_ be a set containing precisely those edges whose terminal vertex is the parent of an isolated leaf. Let *F*_ch_ be a set containing precisely those edges whose terminal vertex is the parent of leaves in an *m*-cherry, except any edge *e* = (*u, υ*) for which some edge in *F*_iso_ lies in the path *P* (*T* ; *ρ, u*). We now combine these two sets, writing *F*_iso_ ∪ *F*_ch_ = *F* = *{f*_1_, *f*_2_, …, *f*_*t*_*}*.

Let *𝓁*^*′*^ : *E*(*T*) → ℝ^*>*0^ be an edge length assignment function on *T* given by *𝓁*^*′*^ (*f*_*k*_) = *M* + *kε* for each *f*_*k*_ ∈ *F* and *𝓁*^*′*^ (*e*) = *ε* for each *e* ∈ *E*(*T*) *\ F*, where 0 *< ε << M*. We ensure that the value of *M* is chosen to be large enough that there are no fixed leaves in 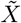 except possibly one fixed leaf whose parent is the root vertex. It is possible to choose *M* with this property by Lemma 8.

Next, we define a number of values based on the *𝓁*^*′*^ edge lengths that will help define the function *𝓁*. For each surviving leaf we calculate the difference between their *φ* index scores, under *𝓁 ′*, before and after the extinction event. That is, for each 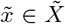 we calculate 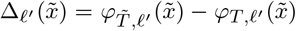. By our choice of *M*, at most one Δ*𝓁′*value is zero and including the *kε* terms in the definition of *𝓁*^*′*^ ensures each Δ*𝓁*^*′*^ value is distinct. Hence we can label the leaves of 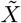 as 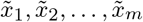 in such a way that 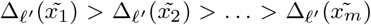. Let 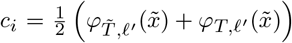 and let *c* = max*{c*_*i*_ : 1 ≤ *i* ≤ *m}*. Finally, let *e*_*i*_ be the pendant edge incident with 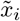 and *P* = *{e*_*i*_ : 1 ≤ *i* ≤ *m}* be the set of these pendant edges.

Now we define the edge length assignment function *𝓁* : *E*(*T*) → ℝ^*>*0^, given by *𝓁*(*e*) = *𝓁* ^*′*^ (*e*) for all *e* ∈ *E*(*T*) *\ P* and *𝓁*(*e*_*i*_) = *c* − *c*_*i*_ + *ε* for all *e*_*i*_ ∈ *P*. Using the edge lengths given by *𝓁*, we see that for each distinct *x*_*i*_ and *x*_*j*_ with *i < j* the following two inequalities hold:

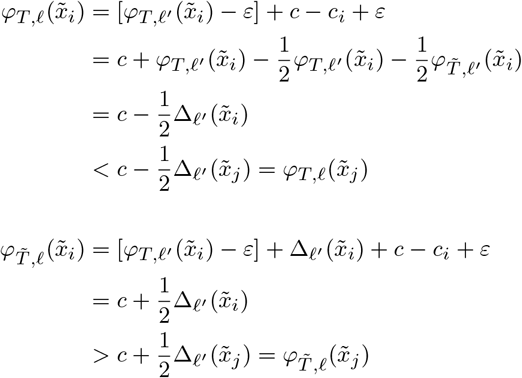

Therefore 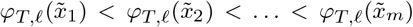 before this extinction event and 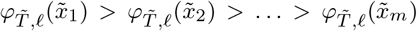 afterwards. That is, there is a strict ranking for the leaves of *T* that is reversible with respect to 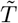. □

We now give a concrete example of the construction used in the proof of Theorem 9. Consider the tree *T* in Figure 3 under the Equal-Splits diversity index. The set *F* in this case consists of three edges: *f*_1_ = (*p, q*), *f*_2_ = (*r, t*) and *f*_3_ = (*r, u*). Suppose that leaves *x*_2_, *x*_5_, *x*_7_, *x*_8_ and *x*_10_ are deleted, inducing the tree 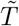 that connects the six remaining leaves. Table 2 gives values for Equal-Splits on both *T* and 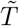 for the surviving species, using an edge length assignment *𝓁 ′* that uses a value of *M* = 32. This table also shows values for 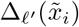 and *c*_*i*_ for each leaf 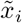. The value of *c* is indicated in bold. The next columns give the lengths of the pendant edges under the assignment function *𝓁* and values for Equal-Splits on both *T* and 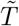 using an edge length assignment *𝓁*. We have chosen to simplify the table by not including the terms of order *O*(*ε*) where they do not impact the rankings we construct. Note that the ordering of the values is opposite in the final two columns, showing that this selection of edge lengths has indeed induced a reversible ranking of ES scores. Figure 7 shows the tree *T* with the edge lengths given by *𝓁* labelled.

**Table 2:** Values used to establish edge lengths for the tree *T* in Figure 7 that lead to a strict and reversible ranking of Equal-Splits index scores. The values are determined in accordance with the construction described in the proof of Theorem 9 with a chosen value of *M* = 32.

| $x_i$ | $ES_{T,\ell'}(x_i)$ | $ES_{\tilde{T},\ell'}(x_i)$ | $\Delta_{\ell'}(x_i)$ | $c_i$ | $\ell(e_i)$ | $ES_{T,\ell}(x_i)$ | $ES_{\tilde{T},\ell}(x_i)$ |
| --- | --- | --- | --- | --- | --- | --- | --- |
| $x_6$ | 8 | 32 | 24 | 20 | 4 | 12 | 36 |
| $x_9$ | 16 | 32 | 16 | **24** | $\varepsilon$ | 16 | 32 |
| $x_4$ | 8 | 16 | 8 | 12 | 12 | 20 | 28 |
| $x_1$ | 2 | 8 | 6 | 5 | 19 | 21 | 27 |
| $x_3$ | 4 | 8 | 4 | 6 | 18 | 22 | 26 |
| $x_{11}$ | $\varepsilon$ | $\varepsilon$ | 0 | $\varepsilon$ | 24 | 24 | 24 |

**Fig. 7:**
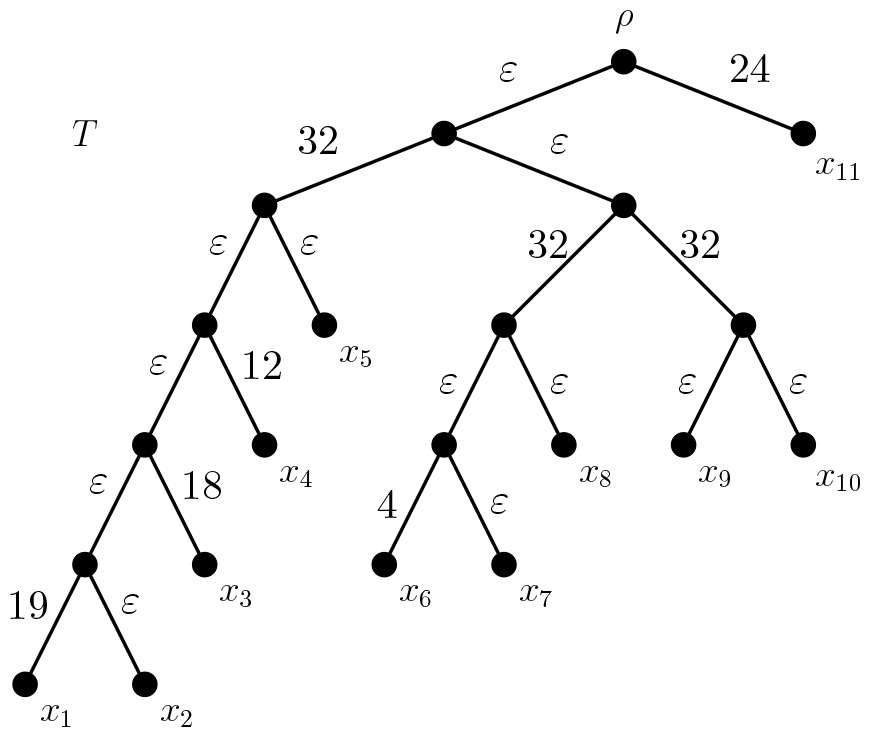
Rooted phylogenetic tree *T* labelled with edge lengths. For these edge lengths, the extinction of leaves *x*_2_, *x*_5_, *x*_7_, *x*_8_ and *x*_10_ leads to a reversal of Equal-Splits rankings for the surviving species.

## 5 Robustness of diversity indices under the ultrametric constraint

We now consider the rankings of diversity index scores on phylogenetic trees whose edge lengths obey the ultrametric constraint. Fischer and colleagues noted that for a rooted ultrametric caterpillar tree no set of extinctions is able to change the ranking order of the remaining leaves [14, Proposition 1]. However, we shall show here that this robustness does not extend to all ultrametric rooted phylogenetic trees, by giving examples of complete ranking reversals for FP and ES. To understand how these example trees were found, we first outline some necessary conditions for the reversibility of FP index scores in the ultrametric context. Note that, to simplify the presentation in this section, we restrict our attention to rooted binary phylogenetic trees.

For every diversity index *φ* the ranking *π*(*X, φ*_*T*_) on an ultrametric rooted binary phylogenetic *X*-tree *T* cannot be strict, as both leaves from any ultrametric cherry must have equal index scores. With this in mind, we impose some further conditions that are not explicitly addressed by the earlier definition of a reversible diversity index. For *x*_*i*_, *x*_*j*_ ∈ *X*, whenever *π*_*T*_ (*x*_*i*_) *< π*_*T*_ (*x*_*j*_) we require 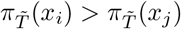 for 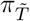 to be considered to be in the opposite order to *π*_*T*_. That is, it is not enough for 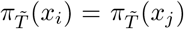. Moreover, to eliminate trivial reversals, for a ranking 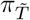 to be considered to be in the opposite order to *π*_*T*_, we require *π*_*T*_ (*x*_*i*_) *< π*_*T*_ (*x*_*j*_) and 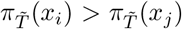 for at least one pair of distinct leaves *x*_*i*_, *x*_*j*_. The necessary conditions for a Fair Proportion ranking reversal under the ultrametric constraint are given in Proposition 10 and Corollary 11. It can be easily seen that caterpillar trees do not meet the criterion below. However many other ultrametric rooted phylogenetic trees do, as shown in Section 5.1.

### Proposition 10

*Let T be a rooted binary phylogenetic X-tree whose edge lengths satisfy the ultrametric condition and let π*_*T*_ *be the ranking of FP index scores for T*. *Suppose x* ∈ *X is a leaf vertex contained in no cherry of T*. *Let υ be the parent vertex of x and write c*_*T*_ (*υ*) *as the disjoint union {x}* ∪ *A, where A contains at least two distinct leaves of T* .

*Then there exist ultrametric edge lengths and a set of extinctions for which the ranking π*_*T*_ *is reversible only if, whenever x survives, the entire set A has become extinct*.

*Proof* Let *x* ∈ *X* be a leaf of *T* that is not contained in any cherry, and let *υ* be the parent vertex of *x*. As *x* is not in a cherry, *c*_*T*_ (*υ*) must contain at least two distinct leaf vertices different from *x*, say *y* and *z*.

Then *FP*_*T*_ (*x*) must be larger than *FP*_*T*_ (*y*). To see this, first note that both *x* and *y* are allocated the same proportion of edge lengths along the path from the root vertex to *υ*. As *T* satisfies the ultrametric condition, the length of edge (*υ, x*) is the same as the length of the path between *υ* and *y*. Yet while leaf *x* is allocated the entire length of edge (*υ, x*), leaf *y* is only allocated part of the total length of the edges along the path from *υ* to *y*. This is because *y* must share some of this total with *z* (from along the edges connecting *υ* to the common parent of *y* and *z*).

Next, suppose a set of extinctions occurs that both *x* and *y* survive and that 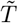 is the resulting phylogenetic tree. Then 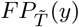 can be no larger than 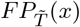, with equality holding if and only if *x* and *y* form a cherry in 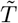. Hence *x* and *y* are not ranked in the opposite order in 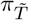 compared to *π*_*T*_. Hence the ranking *π*_*T*_ is not reversible if *y* survives alongside *x*. Repeating this observation for all vertices in *c*_*T*_ (*υ*) (other than *x*) gives the stated result. □

### Corollary 11

*Let T be a rooted binary phylogenetic X-tree whose edge lengths satisfy the ultrametric condition and let π*_*T*_ *be the ranking of FP index scores for T*. *There exist ultrametric edge lengths and a one-per-cherry extinction event for which the ranking π*_*T*_ *is reversible only if every leaf in X is a member of some cherry*.

*Proof* Let *x* ∈ *X* be a leaf of *T* that is not contained in any cherry, and let *υ* be the parent vertex of *x*. As *x* is not in a cherry, *c*_*T*_ (*υ*) must contain at least two distinct leaf vertices different from *x*.

Choose two vertices from *c*_*T*_ (*υ*) that form a cherry, say *y* and *z*. Suppose that, say, *y* survives a one-per-cherry extinction event. Since *x* is not contained in any cherry of *T*, then *x* must also survive the extinction event. Therefore, by Proposition 10, the ranking *π*_*T*_ is not reversible. □

### 5.1 Examples of ultrametric phylogenetic *X*-trees with reversible rankings

We present a family of ultrametric rooted binary phylogenetic trees with reversible rankings under both the Fair Proportion and Equal-Splits indices. Moreover we give specific edge lengths to illustrate such reversals for both indices after the extinction of one leaf per cherry. The family is illustrated by a representative tree *U* in Figure 8a, with twelve leaves: *x*_1_, …, *x*_12_. A similar tree can be constructed for any even number of leaves *n*, where the sum of the edge lengths on the unique path from the root vertex to a leaf is *n* – 1 and the pendant edges right-to-left follow the pattern of increasing positive integer values as shown. A second reversible ultrametric family, consisting of balanced phylogenetic trees, is presented in supplementary material, as well as some further individual examples.

**Fig. 8:**
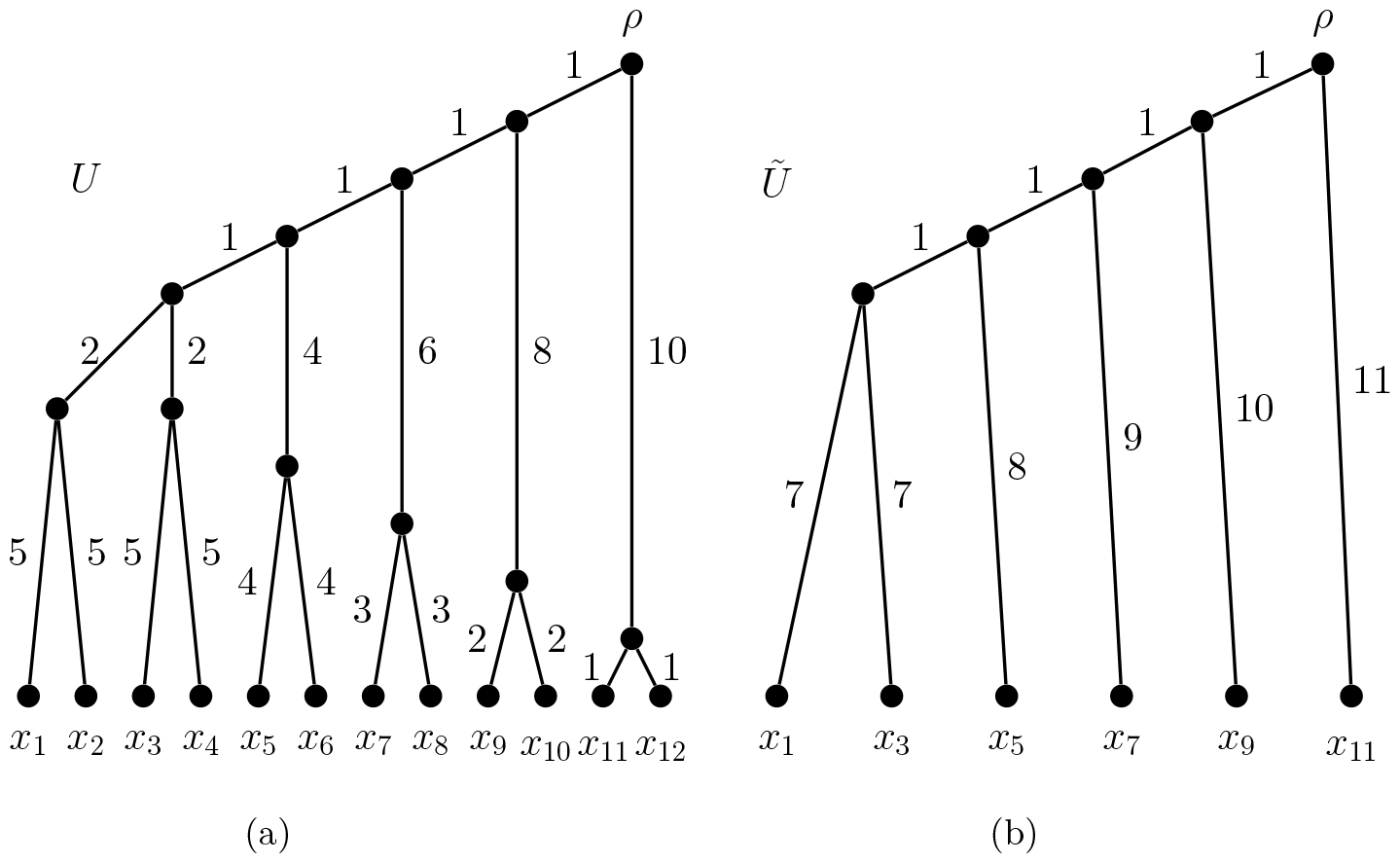
An ultrametric rooted phylogenetic tree *U* before and after a set of extinctions that reverses both the FP and ES index score rankings for surviving species. See Table 3 for details.

Let *U* be the rooted phylogenetic tree in Figure 8a with ultrametric edge lengths as marked. The tree *Ũ* in Figure 8b is obtained from *U* after the extinction of every species with an even subscript. The Fair Proportion index scores for each surviving leaf are given in Table 3. Let *π*_*U*_ and *π*_*Ũ*_ be rankings of FP index scores for the trees *U* and *Ũ* respectively. Observe that *π*_*Ũ*_ ranks the species in the opposite order to *π*_*U*_, hence *π*_*U*_ is reversible. (The scores of leaves *x*_1_ and *x*_3_ remain equal before and after the extinctions.) Table 3 also shows that the same behaviour occurs under the Equal-Splits index on *U*. This example answers the question posed in [14] by showing that the ‘worst-case’ scenario is possible under the ultrametric constraint.

**Table 3:** Fair Proportion and Equal-Splits index values for *U* and *Ũ*, rounded to three decimal places. The largest value in each column appears in boldface.

| $x$ | $FP_U(x)$ | $FP_{\tilde{U}}(x)$ | $ES_U(x)$ | $ES_{\tilde{U}}(x)$ |
| --- | --- | --- | --- | --- |
| $x_1$ | **6.642** | 8.283 | **6.469** | 7.938 |
| $x_3$ | **6.642** | 8.283 | **6.469** | 7.938 |
| $x_5$ | 6.392 | 8.783 | 6.438 | 8.875 |
| $x_7$ | 6.225 | 9.450 | 6.375 | 9.750 |
| $x_9$ | 6.100 | 10.200 | 6.250 | 10.500 |
| $x_{11}$ | 6.000 | **11.000** | 6.000 | **11.000** |

## 6 Concluding remarks

Diversity indices offer us the ability to take a rooted phylogenetic tree that describes the evolution of a *set* of species and quantify the evolutionary history of species individually. Helpfully, the ranking of these diversity index scores can indicate priorities for conservation. However, the usefulness of these rankings is diminished somewhat if they are inconsistent over time. Were the changes to diversity rankings small in nature, they might be easily ignored. But the potential for the complete reversal of these rankings could cause uncertainty over the usefulness of diversity indices in general. Fischer and colleagues [14] do not conclude that reversibility was grounds for disregarding the use of the Fair Proportion index entirely, but that it was an effect that needed to be kept in mind. They suggest that taking various extinction events into account could be an important consideration before applying diversity indices. Based on Theorems 5 and 9 here, we suggest that the lack of robustness is an unavoidable part of using phylogenetic diversity indices, apart from possibly some boundary indices that seem quite unrealistic. These results indicate that the lack of a diversity index that is both robust and biologically reasonable is because such an index does not exist.

As such, reversibility is less a property to be held against FP or ES in preference to other diversity indices and rather more an aspect of measuring at a species-by-species level starting from a phylogenetic tree. We should not be surprised that a species’ contribution to the phylogenetic diversity of a larger set changes (usually increasing) given the demise of close relatives. Thus a diversity index that sensibly measures this contribution cannot be expected to fix its scores in the face of ongoing extinction events. Combinations of extinctions will likely lead to many score changes (of differing magnitudes) that could upset the initial ranking. In the extreme, we have seen complete reversals of rankings are possible.

The examples given in this paper have shown that ranking reversals may be induced not only given quite varied edge lengths but also on rooted phylogenetic trees under the ultrametric constraint. It may be useful for further investigation to determine the minimum number of extinctions required to cause an FP ranking reversal on ultrametric trees. This number will likely depend on the total number of leaves in a tree, as well as the number of leaves allowed to have the same FP index score despite not sharing a cherry. An understanding of these minimal extinction events could help to determine whether the converse of Proposition 10 holds and the extent to which real, time-based phylogenies may be susceptible to reversals.

Our presentation has largely been concerned with the theoretical worst-case scenarios. For this reason, the examples provided may not seem particularly realistic or relevant, especially given the large number of simultaneous extinctions required for the effects shown. The one-per-cherry extinction for binary non-rigid indices, the further isolated leaf extinctions for rigid indices and the even more extinctions required on nonbinary trees all push the limits of plausibility. Furthermore, Proposition 10 demands an even larger number of extinctions before an ultrametric ranking reversal can occur, namely at least half of all species. This is likely an unrealistic number of extinctions to occur together, but the point is that the extreme scenario is theoretically possible. We hope that the negative effects of ranking disruption in real scenarios are brought into focus by the stronger results shown in theory. Indeed, while simultaneous extinctions may seem unlikely, the timescale of progressive extinctions is still possibly shorter than that on which the related conservation efforts are able to adapt and reprioritise. Short of a complete reversal are many types of ranking alterations that would be quite unsettling to a co-ordinated programme of conservation. In addition, Fischer and colleagues investigated the effect of one-per-cherry extinctions on 575 real phylogenies under Fair Proportion and found dramatic re-orderings of FP index scores were indeed possible with real data [14].

Finally, phylogenetic diversity indices, as have been used until now and as were described in [16], have been based on the assumption that, while edge lengths are used to calculate particular score values, the method of calculation should be independent of the lengths themselves. This has not been adequately justified apart from on the grounds of mathematical simplicity and the fact that the two diversity indices used in practice (FP and ES) both have this property. The widespread nature of diversity index ranking reversals suggests there may be value in using a measure that takes into account edge lengths more directly, if doing so could minimise ranking disruptions.

## Acknowledgments

The author was supported by the New Zealand Marsden Fund (MFP-UOC2005). I thank Charles Semple and Mike Steel for their helpful comments and questions and two anonymous referees for their constructive suggestions.

## Declarations

The author has no conflicts of interest to declare that are relevant to the content of this article.

## References

[1] Redding, D.W., Mazel, F., Mooers, A.Ø.: Measuring evolutionary isolation for conservation. PLoS One 9(12), 113490 (2014)

[2] Isaac, N.J., Turvey, S.T., Collen, B., Waterman, C., Baillie, J.E.: Mammals on the edge: conservation priorities based on threat and phylogeny. PloS One 2(3), 296 (2007)

[3] Bordewich, M., Rodrigo, A.G., Semple, C.: Selecting taxa to save or sequence: desirable criteria and a greedy solution. Systematic biology 57(6), 825–834 (2008)

[4] Jetz, W., Thomas, G.H., Joy, J.B., Redding, D.W., Hartmann, K., Mooers, A.O.: Global distribution and conservation of evolutionary distinctness in birds. Current Biology 24, 919–930 (2014)

[5] Forest, F., Moat, J., Baloch, E., Brummitt, N.A., Bachman, S.P., Ickert-Bond, S., Hollingsworth, P.M., Liston, A., Little, D.P., Mathews, S., et al.: Gymnosperms on the edge. Scientific reports 8(1), 1–11 (2018)

[6] EDGE of Existence Programme: EDGE of Existence: Evolutionarily Distinct & Globally Endangered (2022). http://www.edgeofexistence.org Accessed 3 July 2022

[7] Mace, G.M., Gittleman, J.L., Purvis, A.: Preserving the tree of life. Science 300(5626), 1707–1709 (2003)

[8] Davis, M., Faurby, S., Svenning, J.-C.: Mammal diversity will take millions of years to recover from the current biodiversity crisis. Proceedings of the National Academy of Sciences USA 115, 11262–11267 (2018)

[9] White, T.B., Petrovan, S.O., Christie, A.P., Martin, P.A., Sutherland, W.J.: What is the price of conservation? A review of the status quo and recommendations for improving cost reporting. BioScience 72(5), 461–471 (2022)

[10] Griffiths, R., Bell, E., Campbell, J., Cassey, P., Ewen, J., Green, C., Joyce, L., Rayner, M., Toy, R., Towns, D., Wade, L., Walle, R., Veitch, C.: Costs and benefits for biodiversity following rat and cat eradication on Te Hauturu-o-Toi/Little Barrier Island. In: Veitch, C.R., Clout, M.N., Martin, A.R., Russell, J.C., West, C.J. (eds.) Island Invasives: Scaling up to Meet the Challenge. Occasional Paper SSC, pp. 558–567. IUCN, Gland, Switzerland (2019)

[11] Redding, D.: Incorporating genetic distinctness and reserve occupancy into a conservation priorisation approach. Master’s thesis: University of East Anglia (2003)

[12] Shapley, L.S.: A value for n-person games. Annals of Mathematics Studies 28, 307–318 (1953)

[13] Fuchs, M., Jin, E.Y.: Equality of shapley value and fair proportion index in phylogenetic trees. Journal of mathematical biology 71, 1133–1147 (2015)

[14] Fischer, M., Francis, A., Wicke, K.: Phylogenetic diversity rankings in the face of extinctions: The robustness of the fair proportion index. Systematic Biology 72(3), 606–615 (2023)

[15] Bordewich, M., Semple, C.: Quantifying the difference between phylogenetic diversity and diversity indices. arXiv preprint arXiv:2304.10725 (2023)

[16] Manson, K., Steel, M.: Spaces of phylogenetic diversity indices: Combinatorial and geometric properties. Bulletin of Mathematical Biology 85 (2023)

[17] Faith, D.P.: Conservation evaluation and phylogenetic diversity. Biological Conservation 61, 1–10 (1992)

