## Supplementary Material for "The robustness of phylogenetic diversity indices to extinctions"

### The robustness of phylogenetic diversity indices to extinctions: supplementary material

Kerry Manson

School of Mathematics and Statistics, University of Canterbury, Christchurch, New Zealand.

Contributing authors:;

#### 1 Types of diversity measure

##### 1.1 Rigid diversity indices

For small phylogenetic trees we can easily describe all rigid diversity indices. Consider the five-leaf caterpillar tree $Cat_5$ shown in Figure 1, where each edge has unit length. In [1], the space of all diversity index score vectors for this tree, $S(Cat_5, \mathbf{1})$, was determined. Let $\varphi$ be a diversity index for $Cat_5$ and let $(x, y) = (\varphi(x_4), \varphi(x_3))$ describe points in the space $S(Cat_5, \mathbf{1})$. To clarify, each point within the space drawn on the right hand side of Figure 1 represents a distinct diversity index on $Cat_5$.

The rigid diversity indices for $Cat_5$ (with unit-length edges) can be found by setting the ratio of allocations at vertex $s$ to be equal to the ratio of allocations at vertex $t$. Those points satisfying this relationship are exactly described by the equation $y = -x^2 + 4x - 2$, for $1 \leq x \leq 2$. The parabolic curve in Figure 1 connecting the diversity indices $\kappa$ and $\nu$ contains these points. Note that the curve passes through the point representing the Equal-Splits index and does not pass through the point representing the Fair Proportion index. Note also that the rigid indices are relatively rare, lying on a single curve compared to the larger two-dimensional space of all diversity indices.

The rarity of rigid diversity indices extends to other phylogenetic trees beyond this illustrative example. We find rigid diversity indices by taking a pair of isolated vertices with different breadth and setting their ratios of allocations to be equal. For a rooted phylogenetic tree that induces an $m$-dimensional convex space of diversity indices, there are at most $\binom{m}{2}$ ways to choose such pairs. Equating two ratios of allocations leaves an $(m-1)$-dimension space of

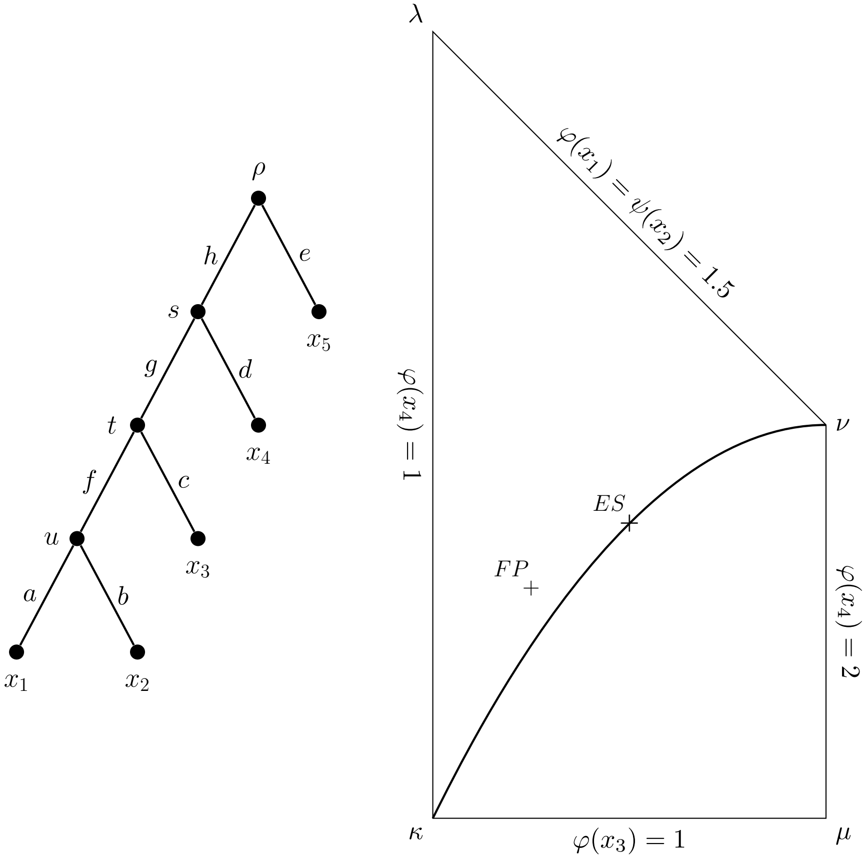


**Fig. 1**: Left: The rooted phylogenetic caterpillar tree $Cat_5$. Right: the space of diversity indices $S(Cat_5, \mathbf{1})$. The parabolic curve that passes through $\kappa$, ES and $\nu$ shows the set of points representing rigid diversity indices. Figure adapted from [1].

feasible solution points, each representing a rigid diversity index. Hence rigid diversity indices lie within a union of at most $\binom{m}{2}$ $(m-1)$-dimensional surfaces within an $m$-dimensional convex space of diversity indices. Although this may represent a large number of points, it will always be outnumbered by the higher-dimensional space of non-rigid diversity indices. Related to this point is that boundary indices are also rare compared to interior indices, being located in one of at most $2m$ $(m-1)$-dimensional spaces on the boundary of $S(T, \ell)$.

#### 1.2 Diversity based on the Banzhaf index

The link between the Fair Proportion index and the Shapley Value leads to the natural supposition that other measures from co-operative game theory may generate useful diversity indices. In particular, we can create a game from a rooted phylogenetic $X$-tree $T$ in the following simple way: the value (or weight) of each set of leaves is the PD score of the minimal subtree connecting those leaves to the root vertex. We write this as $w(S) = PD_T(S)$, where $S$ is a subset of $X$. Then the *Banzhaf value* of a leaf $x \in X$ in this game, $\mathcal{B}(x)$, is given by $\mathcal{B}(x) = \sum_{S \subseteq X} w(S) - w(S \setminus \{x\})$. That is, the Banzhaf value of $x$ is the unweighted sum of $x$'s marginal contributions to the PD scores across every set of leaves.

We show that the Banzhaf value does not always generate a diversity index in the sense of the definition given in [1]. Suppose that the Banzhaf values of the species in a rooted phylogenetic tree are scaled so that they total to the PD score of that tree. For some phylogenetic trees, this results in having leaves whose scaled Banzhaf Value exceeds their maximal possible score under a diversity index. That is, some species are allocated the value of some evolutionary history that they are not descended from.

Specifically, consider the tree $Cat_5$ from Figure 1. When each edge in $Cat_5$ has unit length we get $PD(Cat_5) = 8$. We calculate the Banzhaf value for each leaf and find: $\mathcal{B}(x_1) = \mathcal{B}(x_2) = 30$, $\mathcal{B}(x_3) = 22$, $\mathcal{B}(x_4) = 18$ and $\mathcal{B}(x_5) = 16$. To get the total of these scores to sum to $PD(Cat_5)$ requires dividing each by 14.5. However this results in $x_5$ having an allocation greater than 1, contravening the definition of a diversity index which implies that $\varphi(x_5) = 1$ for every diversity index $\varphi$. From initial investigations, this behaviour is not restricted to the class of caterpillar trees. It appears that the Banzhaf Value tends to 'over-weight' the significance of the pendant edges of a phylogenetic tree and 'under-weight' the interior ones. More investigation of the links between co-operative games and diversity indices could prove fruitful, though we leave that for future work.

### 2 Examples of phylogenetic trees with reversible index rankings

#### 2.1 Small ultrametric trees with reversible Fair Proportion rankings

Figure 2a illustrates a phylogenetic tree $T_1$ that satisfies the ultrametric condition. A reversal of the FP index score ranking may be caused by the extinction of the single leaf $x$ from $T_1$. Before the extinction of $x$, it was the case that $FP_{T_1}(v) = FP_{T_1}(w) = 4 > \frac{23}{6} = FP_{T_1}(y) = FP_{T_1}(z)$. The extinction of $x$ does not change the FP index score of $v$ or $w$, but the FP index score of $y$ and $z$ increase to 4.5.

Let $T_2$ be the rooted phylogenetic tree in Fig. 2b with edge lengths as marked. Suppose that the tree $\tilde{T}_2$ is obtained from $T_2$ after the extinction of $x_3, x_4, x_6$, and $x_8$. The Fair Proportion index scores for each surviving leaf are given in Table 1b. Let $\pi_{T_2}$ and $\pi_{\tilde{T}_2}$ be rankings of FP index scores for the trees $T_2$ and $\tilde{T}_2$ respectively. Observe that $\pi_{\tilde{T}_2}$ ranks the species in the opposite order to $\pi_{T_2}$, hence $\pi_{T_2}$ is reversible.

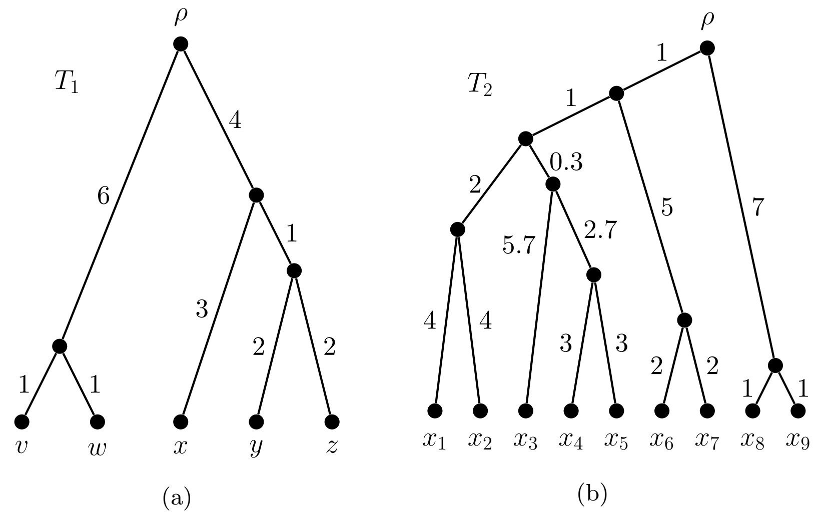


**Fig. 2**: Examples of ultrametric phylogenetic trees whose FP index score rankings are reversible. Reversal occurs with the extinction of $x$ for $T_1$, and the extinction of $\{x_3, x_4, x_6, x_8\}$ for $T_2$.

| $x$ | $FP_{T_3}(x)$ | $FP_{\tilde{T}_3}(x)$ |
|---|---|---|
| $x_1, x_3$ | 14 | **24** |
| $x_5, x_7$ | 14.5 | 23 |
| $x_9, x_{11}$ | 15 | 22 |
| $x_{13}, x_{15}$ | **15.5** | 21 |

(a)

| $x$ | $FP_{T_2}(x)$ | $FP_{\tilde{T}_2}(x)$ |
|---|---|---|
| $x_1, x_2$ | **5.343** | 5.583 |
| $x_5$ | 4.793 | 6.583 |
| $x_7$ | 4.643 | 7.250 |
| $x_9$ | 4.500 | **8.000** |

(b)

**Table 1**: Fair Proportion index values (a) for $T_3$ and $\tilde{T}_3$, and (b) for $T_2$ and $\tilde{T}_2$. The largest value in each column appears in boldface.

#### 2.2 A family of balanced phylogenetic trees with reversible diversity index rankings

Balanced trees are those rooted binary phylogenetic trees where the path from the root vertex to any leaf contains the same number of edges (Figs. 3,4). Let $T_3$ be the rooted phylogenetic tree in Fig. 3 with edge lengths as marked. The tree $\tilde{T}_3$ in Fig. 4 is obtained from $T_3$ after the extinction of every species with an even subscript. The Fair Proportion index scores for each surviving leaf are given in Table 1a. Let $\pi_{T_3}$ and $\pi_{\tilde{T}_3}$ be rankings of FP index scores for the trees

$T_3$ and $\tilde{T}_3$ respectively. Observe that $\pi_{\tilde{T}_3}$ ranks the species in the opposite order to $\pi_{T_3}$, hence $\pi_{T_3}$ is reversible. We note that the FP and ES indices coincide on balanced trees [2]. This example provides an instance of the reversibility of the ES index under the ultrametric condition.

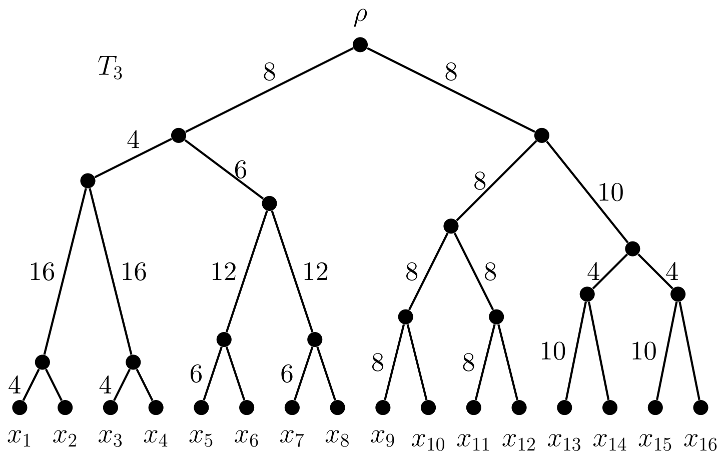


**Fig. 3**: A balanced ultrametric rooted phylogenetic tree with edge lengths marked.

#### 2.3 Specific edge weightings for reversibility

Corollary 1 of [3] states that a strict and reversible ranking exists for any caterpillar tree under the Fair Proportion index. Furthermore, this reversal is caused by the isolated extinction of the species that initially had the lowest FP index score. We now provide a specific assignment of edge lengths on caterpillar trees that exhibits this property.

Consider the tree $Cat_n$ in Fig. 5, and let $\tilde{Cat}_n$ be the tree obtained from $Cat_n$ after the removal of leaf $x_1$, and suppression of the parent vertex of $x_1$ and $x_2$ thereafter. Tables 2 and 3 gives the Fair Proportion index values for the leaves in $Cat_n$ and $\tilde{Cat}_n$ respectively. We include values for an example case where $n = 10$.

We also consider the special case of the reversal of ES index score rankings on the family of caterpillar trees. Consider the rooted caterpillar tree $Cat_n$ with $n \geq 4$, where the pendant edge incident to leaf $x_i$ is labelled $e_i$ and the non-pendant edges are labelled $f_i$ for $1 \leq i \leq n-2$. This tree has one cherry, $\{x_1, x_2\}$, and one isolated leaf $x_{n-1}$. By Theorem 9 edge lengths for $Cat_n$ may be chosen so that the extinction of the two species $x_1$ and $x_{n-1}$ induces an ES ranking reversal. We give a general formula for determining the edge lengths required on such trees. This formula gives an alternative to the construction given in the proof of Theorem 9 and does not use length $\varepsilon$ edges.

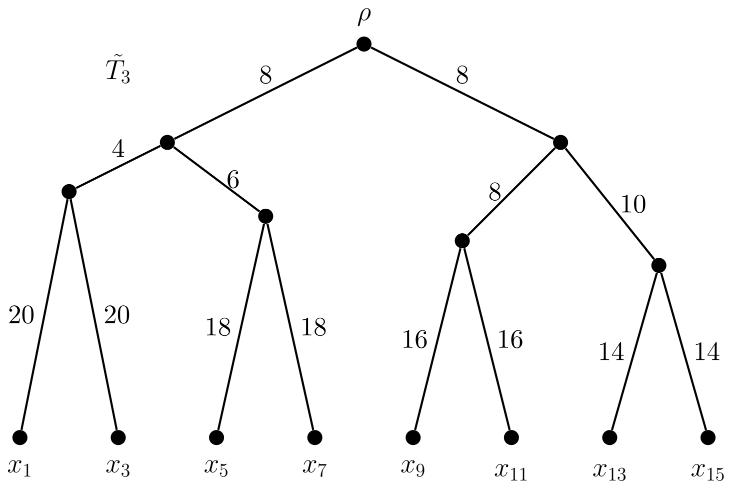


**Fig. 4**: The phylogenetic tree resulting from the extinction of even-indexed species in $T_3$ from Fig. 3. The FP and ES index score ranking of surviving species has been reversed.

Let the edges of $Cat_n$ be assigned lengths by the function $\ell : E(Cat_n) \to \mathbb{R}$ as follows:

- $\ell(e_1) = \ell(e_2) = \ell(e_{n-1}) = 1$
- $\ell(e_i) = 2^{i-2} + n - i - 1$, for $3 \leq i \leq n-2$
- $\ell(e_n) = 2^{n-2} + n - 3$
- $\ell(f_i) = 2^{n-2}$, for all $1 \leq i \leq n-2$

Let $\tilde{Cat}_n$ be the resulting phylogenetic tree once leaves $x_1$ and $x_{n-1}$ are deleted. Table 4 shows the ES index scores for $Cat_n$ and $\tilde{Cat}_n$, illustrating the ranking reversal. In the specific case where $n = 10$, the values for $e_1, e_2, \ldots, e_{10}$ are $1, 1, 8, 9, 12, 19, 34, 65, 1$ and $263$, and $f_i = 256$ for $1 \leq i \leq 8$. The index scores for this scenario appear in Table 4.

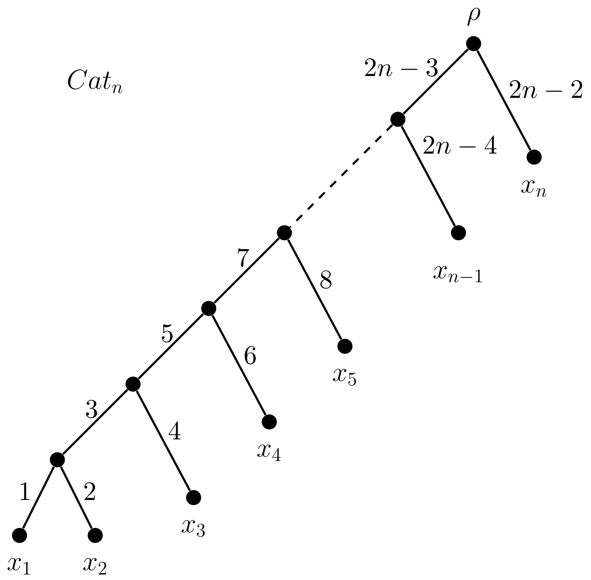


**Fig. 5**: A specific pattern of edge lengths that leads to a reversal of FP index score rankings following the extinction of $x_1$.

| $x$ | $FP_{Cat_n}(x)$ | $n = 10$ |
|---|---|---|
| $x_1$ | $1 + \frac{3}{2} + \frac{5}{3} + \frac{7}{4} + \frac{9}{5} + \cdots + \frac{2n-3}{n-1}$ | 15.171 |
| $x_2$ | $2 + \frac{3}{2} + \frac{5}{3} + \frac{7}{4} + \frac{9}{5} + \cdots + \frac{2n-3}{n-1}$ | 16.171 |
| $x_3$ | $4 + \frac{5}{3} + \frac{7}{4} + \frac{9}{5} + \cdots + \frac{2n-3}{n-1}$ | 16.671 |
| $x_4$ | $6 + \frac{7}{4} + \frac{9}{5} + \cdots + \frac{2n-3}{n-1}$ | 17.004 |
| $x_5$ | $8 + \frac{9}{5} + \cdots + \frac{2n-3}{n-1}$ | 17.254 |
| $\vdots$ | $\vdots$ | $\vdots$ |
| $x_{n-1}$ | $2n - 4 + \frac{2n-3}{n-1}$ | 17.889 |
| $x_n$ | $\mathbf{2n - 2}$ | **18.000** |

**Table 2**: FP index scores for caterpillar tree $Cat_n$. The largest value in the column appears in boldface.

| $x$ | $FP_{\tilde{Cat}_n}(x)$ | $n=10$ |
|---|---|---|
| $x_1$ | extinct | — |
| $x_2$ | $\mathbf{5+\frac{5}{2}+\frac{7}{3}+\frac{9}{4}+\ldots+\frac{2n-3}{n-2}}$ | **20.718** |
| $x_3$ | $4+\frac{5}{2}+\frac{7}{3}+\frac{9}{4}+\cdots+\frac{2n-3}{n-2}$ | 19.718 |
| $x_4$ | $6+\frac{7}{3}+\frac{9}{4}+\cdots+\frac{2n-3}{n-2}$ | 19.218 |
| $x_5$ | $8+\frac{9}{4}+\cdots+\frac{2n-3}{n-2}$ | 18.885 |
| $\vdots$ | $\vdots$ | $\vdots$ |
| $x_{n-1}$ | $2n-4+\frac{2n-3}{n-2}$ | 18.125 |
| $x_n$ | $2n-2$ | 18.000 |

**Table 3**: FP index scores for caterpillar tree $\tilde{Cat}_n$. The largest value in the column appears in boldface.

| $x$ | $ES_{Cat_n}(x)$ | $n=10$ | $ES_{\tilde{Cat}_n}(x)$ | $n=10$ |
|---|---|---|---|---|
| $x_2$ | $2^{n-2}$ | 256 | $\mathbf{2^{n-1}+1}$ | **513** |
| $x_{n-2}$ | $2^{n-2}+1$ | 257 | $2^{n-2}+2^{n-4}+1$ | 321 |
| $x_{n-3}$ | $2^{n-2}+2$ | 258 | $2^{n-2}+2^{n-5}+2$ | 290 |
| $\vdots$ | $\vdots$ | $\vdots$ | $\vdots$ | $\vdots$ |
| $x_5$ | $2^{n-2}+n-6$ | 260 | $2^{n-2}+2^3+n-6$ | 268 |
| $x_4$ | $2^{n-2}+n-5$ | 261 | $2^{n-2}+2^2+n-5$ | 265 |
| $x_3$ | $2^{n-2}+n-4$ | 262 | $2^{n-2}+2^1+n-4$ | 264 |
| $x_n$ | $\mathbf{2^{n-2}+n-3}$ | **263** | $2^{n-2}+n-3$ | 263 |

**Table 4**: Reversal of ES index scores for the $n$-leaf caterpillar tree $Cat_n$ under the extinction of $x_1$ and $x_{n-1}$. Calculated index scores are provided for $n$ in general and the case where $n=10$. The largest value in each column appears in boldface.

### References

[1] Manson, K., Steel, M.: Spaces of phylogenetic diversity indices: Combinatorial and geometric properties. Bulletin of Mathematical Biology **85** (2023)

[2] Wicke, K., Steel, M.: Combinatorial properties of phylogenetic diversity indices. Journal of Mathematical Biology **80**(3), 687–715 (2020)

[3] Fischer, M., Francis, A., Wicke, K.: Phylogenetic diversity rankings in the face of extinctions: The robustness of the fair proportion index. Systematic Biology **72**(3), 606–615 (2023)
